# Deep characterization of cancer drugs mechanism of action by integrating large-scale genetic and drug screens

**DOI:** 10.1101/2022.10.17.512424

**Authors:** Sanju Sinha, Neelam Sinha, Eytan Ruppin

## Abstract

Knowing a drug’s mechanism of action (MOA) is essential for its clinical success by selecting the best indications, likely responders, and combinations. Yet knowledge of many drugs’ MOA remains lacking. Here we present DeepTarget, a computational tool for deep characterization of cancer drugs’ MOA by integrating existing large-scale genetic and drug screens. Spanning ∼1500 drugs across ∼18K possible target genes, DeepTarget predicts: (1) a drug’s primary target(s), (2) whether it specifically targets the wild-type or mutated target forms, and (3) the secondary target(s) that mediate its response when the primary target is not expressed. We first tested and successfully validated DeepTarget in a total of eleven unseen gold-standard datasets, with an average AUC of 0.82, 0.77, and 0.92 for the above three prediction abilities, respectively. We then proceed to use it in a wide range of applications: First, we find that DeepTarget’s predicted specificity of a drug to its target is strongly associated with its success in clinical trials and is higher in its FDA-approved indications. Second, DeepTarget predicts candidate drugs for targeting currently undruggable cancer oncogenes and their mutant forms. Finally, DeepTarget predicts new targets for drugs with unknown MOA and new secondary targets of approved drugs. Taken together, DeepTarget is a new computational framework for accelerating drug prioritization and its target discovery by leveraging large-scale genetic and drug screens.

## Introduction

Knowing a drug’s targets and mechanism of action (MOA) is a highly important determinant of its clinical success. It serves as a biomarker to identify its best indications, stratify patients for therapy, and identify drug combinations. In the last few years, multiple integrated experimental and computational approaches have aimed to address this challenge and predict such targets [1-6]. A few notable examples are tepoxalin, specifically killing cells with high ABCB1 expression [1], and vanadium-containing compounds, specifically killing cells with high SLC26A2 expression [1]. Accordingly, many computation tools have recently been developed to apply machine learning to identify a drug’s MOAs by analyzing information about the compound structure, the drug’s pre/post-treatment expression profiles, adverse effects, and binding profiles [4-6]. However, our understanding of their MOA remains poor, as reflected by their poor clinical and approval success rate. Here, we present DeepTarget, a computational approach to do a deep characterization of cancer drugs’ targets and MOA, which relies on integrating large genetic and drug screens in cancer cell lines. Our pipeline is based on the reasoning that the CRISPR-Cas9 knockout (CRISPR-KO) of a gene encoding the protein target of a given drug can mimic the inhibitory effects of that drug, and thus a target can be identified by identifying a CRISPR knockout phenocopied by the drug.

DeepTarget is not the first approach for building drug target predictors by integrating large genetic and drug screens in cancer cell lines. A recent previous study has taken this route and compared drugs and the CRISPR screens to identify the primary target of drugs, studying 400 drugs, revealing that the MCL1 inhibitors may mediate their efficacy via mitochondrial E3 ubiquitin-protein ligases [3]. DeepTarget adopts a same principle that the CRISPR-Cas9 knockout drug target gene can mimic the drug treatment but aims to markedly go beyond previous work in three important dimensions: (1) First, beyond primary target identification, it performs a deeper characterization of the drug’s MOA by predicting whether a drug specifically targets the WT vs. a mutant form of its known target protein and drug’s secondary target(s) that mediate its response in specific cancer indications when the primary target is not expressed. (2) Importantly, we extensively tested and successfully validated DeepTarget in a wide range of independent gold-standard datasets that are not used in training. (3) Studying a host of new applications of DeepTarget, including predicting the likely clinical success and optimal indications of a drug with high accuracy, repurposing existing compounds to target undruggable oncogenes, and identifying unrecognized targets for drugs with unknown MOA. Accordingly, we provide an overview of DeepTarget (section A), then describe a wide range of validation tests of its prediction abilities (Section B). Finally, we present five applications addressing important open challenges in cancer drug discovery (Section C).

## Results

### A. Overview of *DeepTarget* pipeline

The DeepTarget pipeline is based on the hypothesis that the CRISPR-Cas9 knockout (CRISPR-KO) of a gene encoding the protein target of a given drug can mimic the inhibitory effects of that drug treatment. That is, the sensitivity profile observed across different cancer cell lines after the CRISPR-KO’s of the target gene should be similar to that observed after treatment with the said drug and significantly more similar than the sensitivity profiles observed after the KO of the other genes (**Figure 1A, S1**). Based on the above notion, DeepTarget has three main steps:

**Figure 1:**
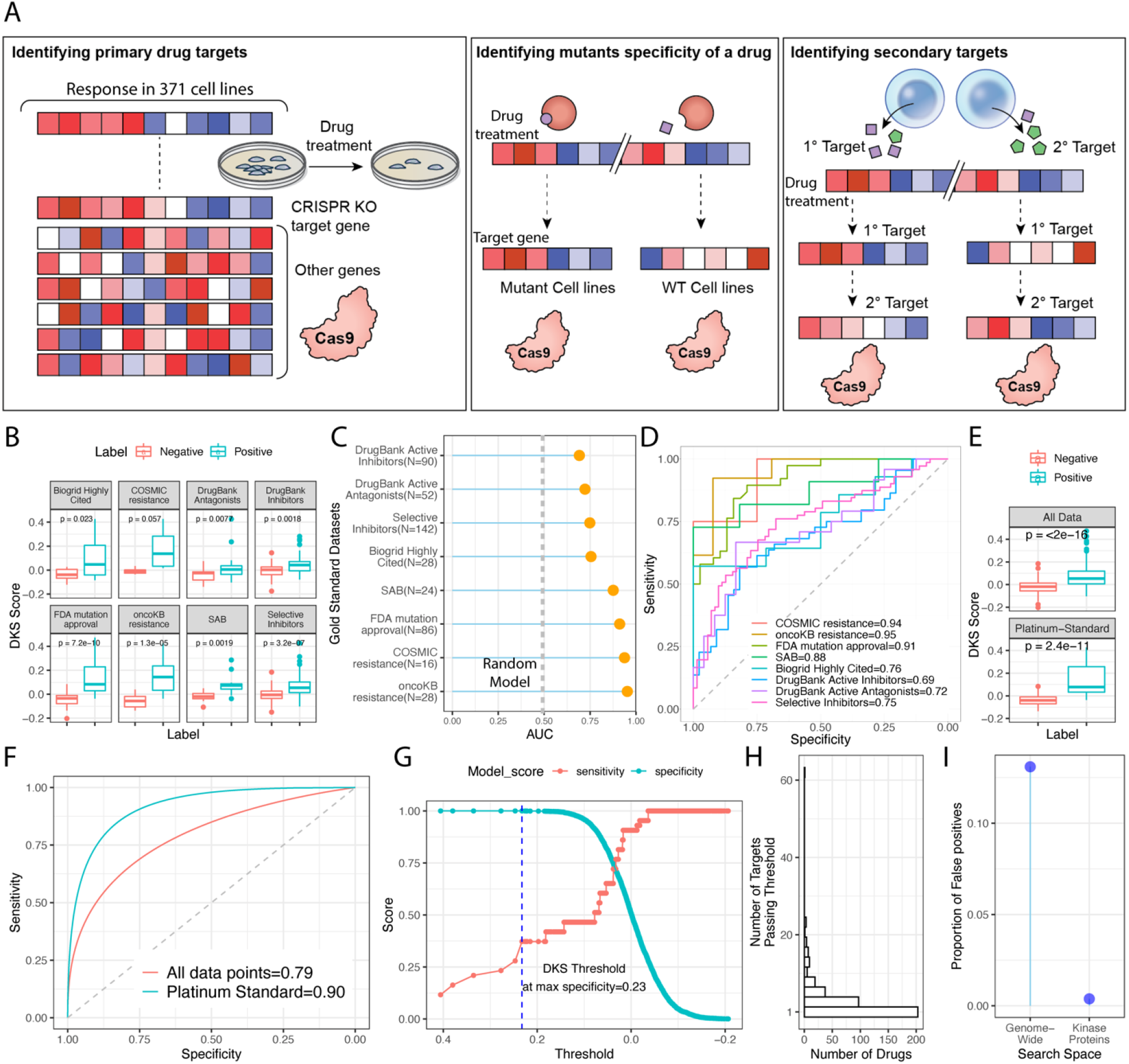
Overview of DeepTarget and its performance across eight gold-standard datasets. **(A)** Overview of the three steps of DeepTarget. (**B**) The DeepTarget predicted DKS score (Y-axis) of high-confidence drug-target pairs (positive, red) vs. negative labels created by shuffling labels (green) from our pipeline across eight gold-standard datasets. Two-sided Wilcoxon rank-sum significance is provided. **(C)** DeepTarget prediction performance was computed using the area under the curve of ROC (X-axis) of our pipeline to identify high-confidence drug-targets pairs from negative labels created using shuffled across eight datasets (Y-axis). The number of data points in each dataset is provided in the dataset name labels on Y-axis. **(D)** The ROC curve for each dataset is provided, showing specificity vs. sensitivity. The respective AUC value is provided at the bottom right. **(E)** DeepTarget predicted DKS score (Y-axis) of platinum-standard drug-target pairs (positive, red) vs. negative labels (green) for all data points, where the ROC curve is provided in **(F). (G)** The relationship between the DKS score threshold of choice and specificity & sensitivity observed in the platinum-standard dataset is shown. The maximum specificity was achieved at the DKS score threshold >0.23. The blue dashed line making this threshold is provided. **(H)** The overall distribution of DeepTarget’s predicted number of targets per drug (X-axis) using this DKS score threshold. **(I)** The proportion of false positives out of total genes (Y-axis) in case of genome-wide search space vs. kinase proteins only.

<u>In the first step, analogous to the approach presented in</u> [3], <u>DeepTarget performs a genome-wide search to predict a drug’s primary target (</u>**<u>Figure 1A</u>-*<u>left panel</u>***). The similarity between the viability profile after CRISPR-KO of a gene and drug treatment is the likelihood for the gene to be the drug’s primary target. To this end, we first mined the large-scale drug response screen (PRISM) [1] for 1450 drugs screens across eight dosages and genome-wide CRISPR-Cas9 knockout viability profiles (CRISPR-KO essentiality, AVANA) from DepMap [7], performed across 371 cancer cell lines present in both the screens (the input to DeepTarget). Integrating these two screens, we calculated a (drug, target) pair-wise similarity score, a Pearson’s correlation Rho, between viability after drug treatment and that observed for each gene CRISPR-KO across 371 cell lines and termed it the ***D****rug-****K****O* ***S****imilarity score (****<u>DKS score</u>***). This score is DeepTarget’s first output and can range from -1 to 1, where 1 signifies a perfect correlation between viability after drug treatment and gene CRISPR-KO. A UMAP based on this DKS score of 1450 drugs across 17386 possible target genes is provided in **Figure S2A-B**. We finally test and validate DeepTarget’s ability to identify primary targets in eight independent datasets (see next section).

<u>In the second step, we predict whether the drug more specifically targets the wild-type (WT) or one of the mutant forms, if any, of its known target protein (</u>**<u>Figure 1A-</u>*<u>middle panel</u>***). We reason that if a drug more specifically targets a mutant form of protein, in the cell lines with this mutant form, the similarity between viability after drug treatment and target CRISPR-KO (their *DKS score*) would be significantly higher than in the cell lines with WT protein. We mathematically model this dependency of the DKS score on the mutation status of the target by employing a regression between the drug and the target’s response across the cell lines, including the mutation status as an interaction term. We call this score the *mutant-specificity* score (this composes the second output of DeepTarget), where a positive mutant-specificity score indicates that a drug differentially targets the mutant form of the target protein more than WT and vice-versa for the negative score. We only model mutation variants present in at least five cell lines providing a minimum statistical power. Given sufficiently large data, we note that different mutation variants can be modeled individually to compute variant-specific targeting.

<u>In DeepTarget’s third and final step, we predict a drug’s secondary target(s)</u> if/when those exist (Figure 1A-right panel). This step is based on the working hypothesis that the secondary target would mediate a drug’s MOA in cell lines where the known primary target is not expressed. To identify such secondary targets, we rank all the genes based on their *DKS score* computed only across the cell lines where the primary target is not expressed, and the resulting top-ranked gene is the predicted secondary target (this secondary-DKS score composes the third output of DeepTarget).

We note that CRISPR-KO of genes downstream or upstream of the actual drug target may also induce a similar sensitivity profile as the actual target and thus may rank high in our primary and secondary target identification steps. Thus, the top-ranked candidates, in addition to a target, also point to other genes involved in the drug’s MOA pathway. This effect will confound DeepTarget’s identification of actual targets, and thus, in a part of our analysis, we additionally introduce a post-filtering step, restricting our target search space to the genes that a drug would likely bind to and thus more likely to be the actual direct targets, e.g., kinase proteins for kinase inhibitors. Finally, when relevant, we also compute the pathways enriched among the top-ranked genes to provide a pathway-level description of each drug’s predicted primary and secondary MOA.

### B. Testing and validating *DeepTarget* predictions

#### Testing *DeepTarget’s* primary target prediction accuracy

After running DeepTarget for the PRISM screen’s drugs and genome-wide CRISPR-KO screen from DepMap as described above, we comprehensively tested its drug-target predictions performance in eight independent gold standard datasets comprising high-confidence drug-target pairs collected from diverse sources (**Table S1**, curation process in **Supplementary Notes 1A**). These datasets include drug-target pairs that have: (1) A mutation in the target gene causing clinical resistance to the drug from COSMIC (*COSMIC resistance, N=16*) [8], or (2) oncoKB (*oncoKB resistance, N=28*) [9], (3) FDA Approval for a target mutation (FDA mutation-approval, N=86) [9], (4) High-confidence as per the scientific advisory board of ChemicalProbes.org (*SAB, N=24*) [10], (5) Multiple independent reports as per BioGrid (Biogrid Highly Cited, N=28) [11], (6) pharmacologically active status as per DrugBank, i.e., the drugs interact directly with the target as part of its MOA and are inhibitors (*DrugBank Active Inhibitors, N=90*) [12] or (7) are antagonists (*DrugBank Active Antagonists, N=52*) [12], (8) Highly selective inhibitors based on their binding profile (SelleckChem selective inhibitors, N=142) [13]. We term the drug-target pairs in any of these resources (some may appear in more than one) as *gold-standard drug-target pairs*, and they serve as our positive labels. We also created two types of negative labels (controls) for each gold-standard dataset by (a) shuffling the target labels (**Methods**) and (b) choosing random protein-coding genes as target labels.

We find that the *gold-standard* drug-target pairs are significantly ranked higher by DeepTarget (i.e., they have a higher *DKS score)* than the negative controls and that DeepTarget stratifies them with a mean AUC of 0.82 across each of the eight different datasets (**Figure 1B-D**). When tested on all data points merged, DeepTarget yields an AUC of 0.79. Looking at the AUCs obtained in each dataset, the AUC for *COSMIC resistance* is 0.94, *oncoKB resistance* is 0.96, FDA mutation-approval is 0.91, SAB is 0.88, Biogrid Highly Cited is 0.76, DrugBank Active Inhibitors is 0.69, DrugBank Active Antagonists is 0.72 and SelleckChem selective inhibitors is 0.75 (**Figure 1B**). Interestingly, DeepTarget’s performance is higher in gold-standard datasets derived from clinical sources vs. in vitro-only sources. We also note that the first three datasets have a confidence score for each drug-target pair, and focusing on pairs with high confidence scores in these datasets yields a mean AUC of 0.96 (**Supplementary Notes 1B**). Furthermore, focusing on DeepTarget predictions for a set of drug-target pairs that are present in more than one gold-standard dataset (termed *platinum-standard*, N=57 pairs) yields an AUC of 0.90 (**Figure 1E-F)**, and quite strikingly, the predictions on the 18 pairs that are present in more than two datasets yield an AUC of 1.

Analyzing this platinum-standard dataset, we found that the threshold of DeepTarget’s DKS score > 0.23 maximizes the specificity of our target identification process (**Figure 1G, Figure S2C**), thus minimizing the likelihood of false positives. Here onwards, we fix this threshold as predictive of whether a gene is a primary target of a drug throughout our study and keep it on value *without any change* for all the validation tests and applications shown in this study, without any further training or modification. This DKS score threshold yielded at least one target for 386 drugs (Total drugs=1450) with an average of 3 targets per drug, where 203 drugs are predicted to have a single target (**Figure 1H**). The overall distribution of the predicted number of targets per drug using this threshold is provided in **Figure 1H**. The above’s stratification power analysis for each drug target class and major pathways separately is provided in **Supplementary Notes 2A**.

We next aimed to mitigate the prediction of highly ranked genes with a similar KO profile to the target as they are downstream or upstream of it but do not bind to the drug (false positives). To this end, we performed the target identification process for drugs with gold-standard target information, but instead of using a genome-wide search space, we limited the target search space to the protein class the drug is designed towards and would thus likely bind to, e.g., kinase proteins for kinase inhibitors. Here, false positives are hits ranked higher than the reported gold-standard target. While restricting our search to the kinase space (N=509 genes) yielded no change in our stratification power (AUC) for identifying the gold-standard targets for kinase inhibitors (0.82); however, the total false positives proportion massively reduced from 13% in the case of genome-wide search to 0.003% (**Figure 1I, Supplementary Notes 2B**, empirical P< 1E-04, computed using random gene sets of the same size). A similar analysis for other major target classes, including GPCRs, ion channels, metabolic enzymes, and nuclear receptors, is provided in **Supplementary Notes 2B**. We thus note that in cases where drug-binding information is available, this strategy could be further employed beyond a genome-wide search.

#### Testing *DeepTarget*’s ability to predict whether a drug specifically targets the wild-type or the mutant form of its known target protein

The second step of *DeepTarget* aims to identify whether a given drug more specifically targets a mutant or the wild-type (WT) form of its known target protein. We hypothesize that if the correlation between drug response and target CRISPR-KO (*DKS Score*) is significantly higher in cell lines harboring a mutant form than in cell lines harboring the WT form of the target, the drug more specifically inhibits the mutant form of the target, and vice-*versa (***Figure 1A-*middle panel***). We performed this for mutation variants present in at least five cell lines. *We modeled* and tested this dependency using a linear regression model between the drug response and target CRISPR-KO efficacies. It includes an interaction term whose coefficient represents how specific a drug may target its mutant vs. WT form (*mutant-specificity score)*. As an initial case study, **Figure 2A** exemplifies DeepTarget’s ability to identify the known mutant-specific targeting for the well-characterized FDA-approved BRAF V600E inhibitor Dabrafenib. Indeed, as hypothesized, Dabrafenib response is significantly correlated with viability after BRAF KO only in cell lines with BRAF V600E mutant.

**Figure 2:**
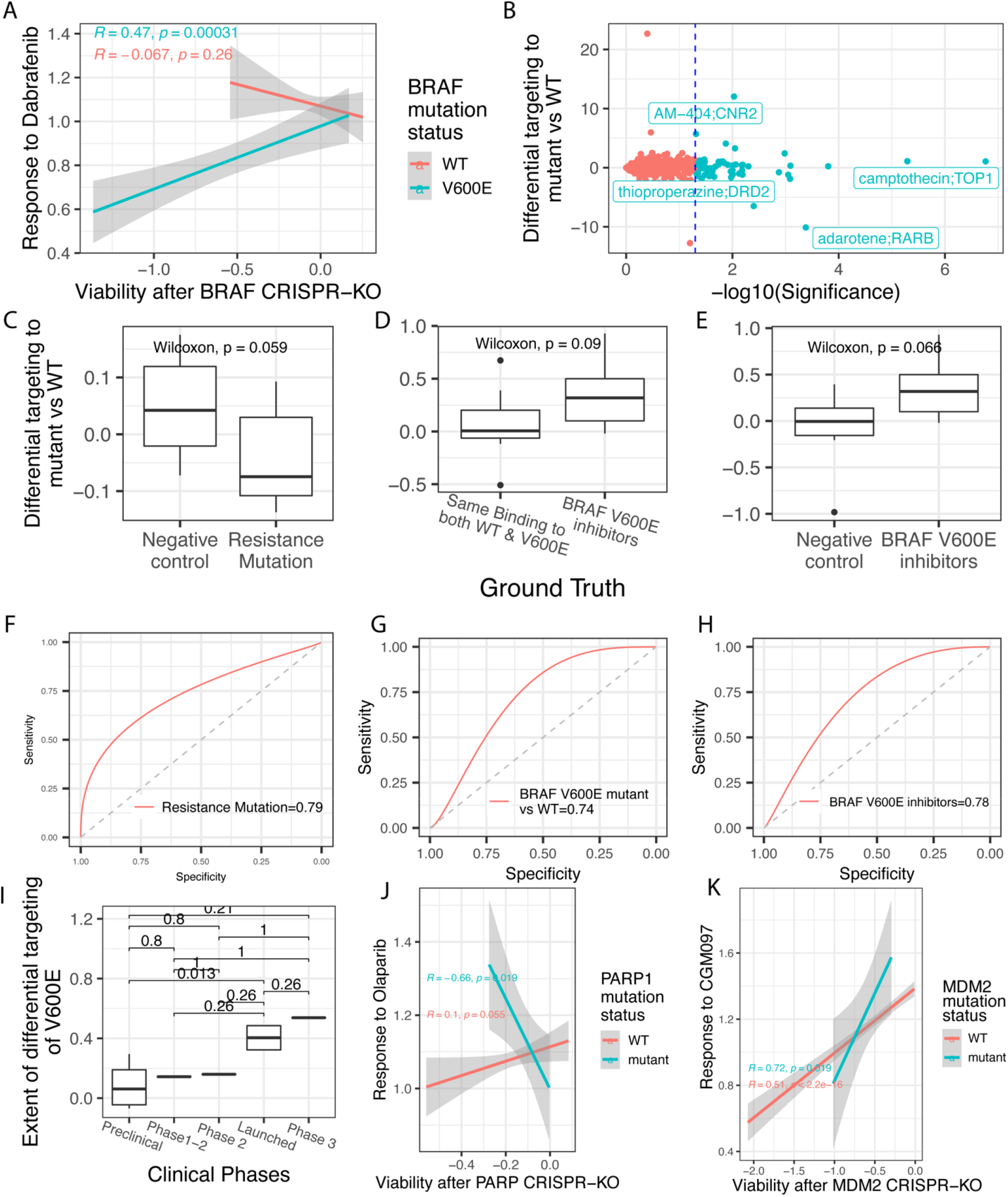
Identifying whether a drug binds more specifically to its primary target protein’s wild-type or mutant form. **(A)** Proof of concept of our pipeline’s ability to predict mutation-specific targeting for Dabrafenib, where the response to dabrafenib is specifically correlated with BRAF in only cell lines with BRAF V600E mutation (green) and not WT (red). The strength of the correlation is provided on the left top. **(B)** The extent of differential binding to the mutant form of the target for all the drugs in our screens (N=1450, y-axis) and the respective significance (X-axis). The statistically significant points are highlighted in green, and the top hits are labeled. **(C)** DeepTarget predicted differential targeting of mutant vs. WT is significantly lower in drugs and clinical resistance mutations from COSMIC vs. shuffled negative labels. The statistical significance is computed using Wilcoxon rank-sum test. **(D)** This is repeated for drugs that bind more specifically to all the drugs in our screens that bind more specifically to BRAF V600E mutation vs. BRAF inhibitors that do not. **(E)** We also compared our prediction vs. negative shuffled labels. **(F-H)** The stratification power of positive and negative labels in **(C-F)** is provided as the area under the ROC curve. **(I)** The extent of mutation-specific targeting (Y-axis) for various clinical phases (X-axis) of BRAF inhibitors is provided. **(J)** Response to Olaparib (Y-axis) vs. viability after PARP1 KO (x-axis) cell lines with PARP mutation (green) vs. WT (red). **(K)** is shown in the same manner as **(J)** for CGM097 and MDM2.

We next computed the *mutant-specificity score* (**Methods**), which quantifies the difference in DKS score between the mutant vs. WT cell lines for all the drugs in our screens (**Figure 2B**, N=1450) and provide this in **Table S2** as a resource. We then tested these predictions in two gold-standard drug sets: The first drug set comprises drugs that have a mutation in the primary target leading to clinical resistance, as reported in oncoKB [9]. These drugs hence by definition, bind less to this resistant mutant form of the target than the WT form, e.g., Y1230H mutation in MET inducing resistance to Crizotinib. Aligned with our hypothesis, we first observed that the mutant-specificity scores of these drugs and the respective target mutant-form pairs are negative, indicating less binding to mutant vs. WT. These scores are also significantly lower for these pairs vs. the negative controls (**Figure 2C, N=16**, Wilcoxon Rank Sum P<0.05). This score can stratify the drugs binding more to WT vs. the mutant form with an AUC of 0.79 (**Figure 2F**). The second gold-standard dataset is comprised of drugs that bind more specifically to the mutant form of the target (curated using binding profiles from SelleckChem, 14). This yielded six drugs, all targeting the BRAF V600E mutation. As negative labels, we found 7 BRAF inhibitors known to bind to V600E and WT form comparably [13]. The predicted V600E *mutant-specificity score* for these drugs is higher for V600E-specific inhibitors vs. ones that bind comparably to V600E and WT (**Figure 2D**). It can stratify mutant-specific drugs with an AUC of 0.74 (**Figure 2G**). As another control, negative shuffled labels also have a significantly lower *mutant-specificity score* (P<0.06, AUC=0.78, **Figure 2E, H**). Finally, we note that the V600E *mutant-specificity score* of BRAF inhibitors can predict how advanced in the clinic a BRAF inhibitor has reached; the higher the *mutant-specificity*, the further in clinical trial stages a BRAF inhibitor reaches (Cor Rho=0.77, P=002, **Figure 2I**). We cautiously note that this analysis may be confounded by the total time since a drug was discovered and introduced in trials (see Methods 6.0 to see how we corrected this confounding factor).

Going beyond BRAF, an approved top example identified by our pipeline is olaparib targeting the PARP1 WT form only. As expected, olaparib response and PARP1 KO are significantly correlated in PARP1 WT cells only (**Figure 2J**). Another notable prediction is that CGM097 specifically targets the *MDM2* mutant form (**Figure 2K**). Given these validations, we applied this analysis to predict six new mutation-specific inhibitors with high clinical advancement and provided these predictions as a resource (**Table S2, N=15**). This includes six drugs specifically targeting mutant form and likely a good candidate due to high specificity to mutant form. It also includes nine drugs where drugs target WT more specifically, and thus, the mutant form will likely induce treatment resistance. The topmost frequent targets of drugs specifically targeting mutant form are BRAF & MDM2, whereas frequent targets of the drugs specifically targeting WT include MAP2K1, FGFR1, and PIK3CA.

#### Testing *DeepTarget*’s ability to predict secondary targets of drugs in indications where the primary target is not expressed

We next test the ability of DeepTarget to predict novel secondary target/s that mediate drug efficacy when the primary target is not expressed, applying the third step of DeepTarget. This analysis is highly relevant for drugs initially designed to target a gene only expressed in a single or a few cancer types. An interesting case in hand is that of Ibrutinib, an FDA-approved drug for multiple hematological malignancies known to target Bruton’s tyrosine kinase (BTK) irreversibly. Even though BTK is specifically expressed in hematological malignancies (**Figure S3A**), Ibrutinib shows anti-tumor efficacy in multiple other solid cancer types (**Figure S3B**). Using *DeepTarget, we* confirm that Ibrutinib’s primary target is likely to be BTK in cell lines with BTK expression (DKS score=0.8, P=0.1, **Figure 3A, highlighted in red, Methods**). However, in contrast, in cell lines with low-BTK expression, there is no correlation between BTK KO viability and Ibrutinib response (DKS score=0, P=0.96, **Figure 3A, highlighted in green**). Focusing on these cell lines with no BTK expression, *DeepTarget* identifies EGFR as Ibrutinib’s alternative secondary target (Rank=1, EGFR DKS score=0.44, BTK DKS score=0.01, **Figure 3B**). Ibrutinib’s kinase binding profile revealed a strong binding to both EGFR and EGFR mutant forms, further supporting this finding [14]. We further note that the other two known BTK inhibitors in our screen, AVL-292, and CNX-774, also show this ability to target EGFR with low BTK activity.

**Figure 3:**
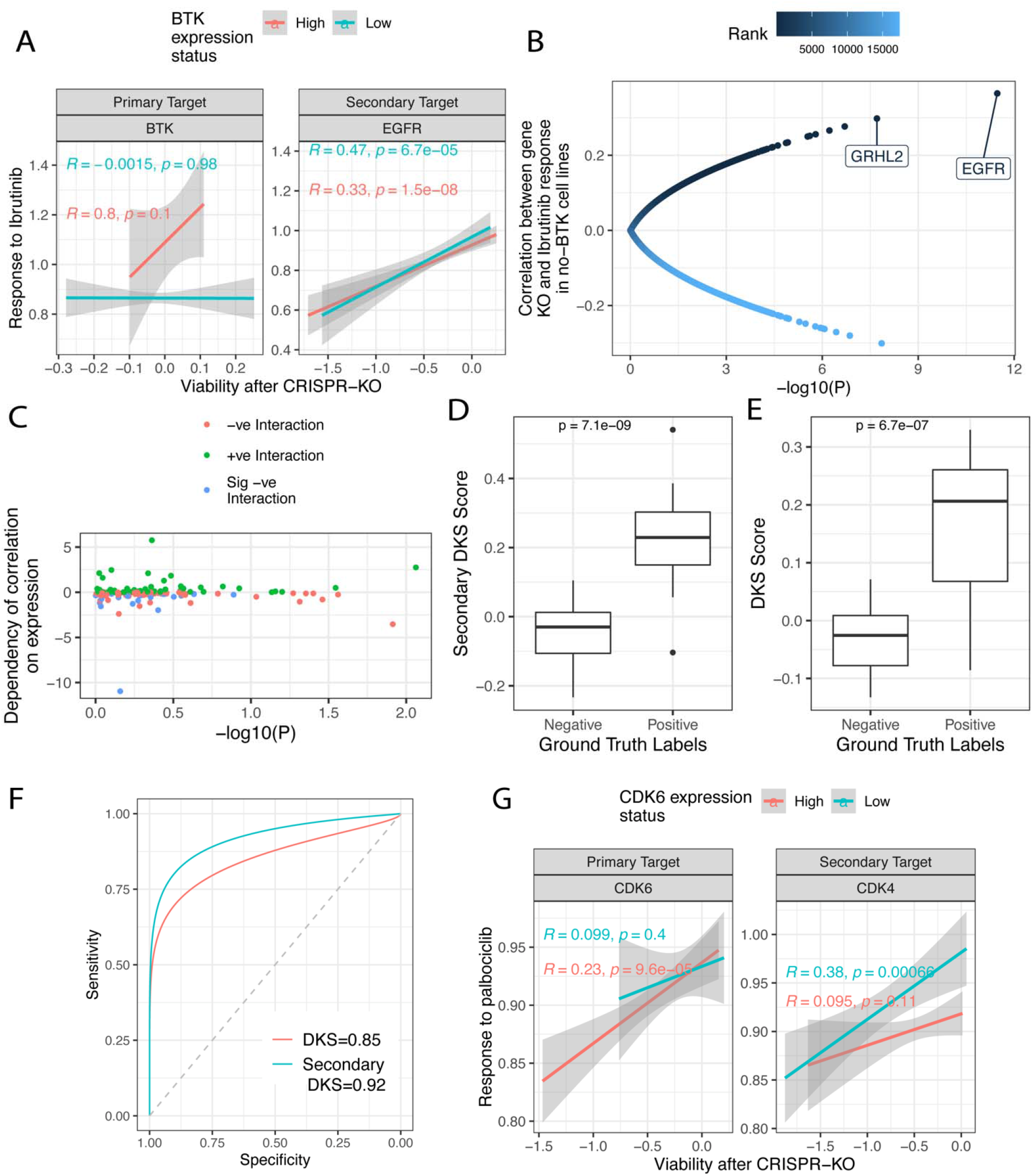
Identifying secondary targets of drugs that mediate the response when the primary target is not expressed. **(A)** Viability after BTK KO (X-axis) vs. Ibrutinib response (Y-axis) in BTK low expression (green, Step 3 score) vs. non-low cell lines (red). The Pearson correlation is provided on the left top with the respective color. **(B)** The strength of the correlation of ibrutinib response and viability after KO of each gene (Y-axis) is provided with respective significance (X-axis) in low-BTK expression cell lines, aimed at identifying the secondary target of Ibrutinib when BTK is not expressed. Each point representing a gene is colored by its predicted rank from our pipeline. The top two hits, EGFR and GRHL2, are labeled. **(C)** The dependency of the DKS score upon the expression levels of the primary target is computed using an interaction term between these two variables in a regression model; the strength (Y-axis) and respective significance are provided. A negative estimate represents that this correlation is decreased in cell lines with low expression of the target. The points are colored by their direction and the significance of this dependency. **(D)** The predicted secondary DKS score of secondary targets from the analysis only using cell lines with low expression of the primary target (Y-axis) of drugs with high-confidence annotation vs. negative labels (X-axis). The DKS score using all cell lines is also provided in **(E)** for comparison. **(F)** The ROC curve represents the prediction accuracy of secondary targets vs. negative labels for Step 1 vs Step 3. The respective area under the ROC curve is provided at the bottom right. **(G)** As an illustration, we show the response to Palbociclib vs. viability after CRISPR KO of CDK6 and CDK4 in low vs. high expression of CDK6 cell lines.

Motivated by this observation, we systematically identified the secondary target/s of all drugs in the screens. To this end, we repeated our target identification process in cell lines where their primary targets are not expressed to produce a *secondary-DKS Score* for such drugs. To this end, we first tested if, indeed, the correlation between the drug response and viability after the primary target CRISPR-KO (*DKS score*) decreases in cell lines where the primary target is not expressed. We found such a reduction in *DKS score* for 76% of the drug-known target pairs that passed DeepTarget’s step 1 (**Figure 3C**). For these drug-target pairs, we next aimed to predict and further test their secondary targets.

To test *DeepTarget* predictions of secondary targets, we assembled a pertaining gold standard dataset. We collected the targets of all drugs with multiple known targets present in at least one of our eight previously described gold standards, whose primary target is ranked as the best predicted in the first step of DeepTarget (N=64). We then used their secondary-target annotations in these datasets as a gold standard benchmark for testing and validating the predictions made in this step. *DeepTarget identified the* secondary targets of these 64 drugs vs. negative labels with an AUC of 0.92 (**Figure 3D, 3F, S4**). We note, however, that performing this prediction based on step 1 based DKS Score (correlation based on *all cell lines*) also identifies the secondary targets with a quite high AUC of 0.85 (**Figure 3E, 3F**), which is, however, still significantly less than that obtained using the secondary target identification step (Wilcoxon rank-sum P= 2E-04).

One notable top hit arising in this analysis is palbociclib, an FDA-approved treatment for breast cancer that selectively inhibits cyclin-dependent kinases CDK4 and CDK6. As evident in **Figure 3G**, in cell lines with CDK6 expression, it is indeed the primary target, but when CDK6 is lowly expressed, CDK4 becomes the main target. Interestingly, Abemaciclib and Ribociclib, the other approved inhibitors of CDK4/6, do not show this hierarchical targeting. We provided numerous such examples where DeepTarget correctly predicted the known primary and secondary targets of drugs in **Supplementary Notes 3, Figure S4-5**. A notable frequent secondary target is EBBR3, predicted to be a secondary target of multiple known EGFR inhibitors in cell lines with low EGFR activity.

### C. Applications of *DeepTarget*

#### The predicted drug’s specificity to its target is strongly associated with its clinical advancement

We hypothesized that the specificity of a drug to bind and inhibit its target might predict its clinical advancement, where the drugs highly specific to their target have a higher chance of advancing in clinical trials. We proposed that DeepTarget’s DKS score of a drug for its known target would indicate this specificity (termed *drug-target specificity*) and thus can subsequently determine its clinical advancement (more details in **Methods section 6.0**). The term *clinical advancement* here refers to the stage a given drug has reached, starting from the preclinical stage to various clinical trial phases (1 to 3) and, finally, its approval. As a first test case, we tested this hypothesis for EGFR, the target with the most drugs across all different phases of clinical trials. We have had 41 drugs in our screens that are known to target EGFR. Among these, 8 are FDA approved, 6 are in each Phase (1, 2 & 3), and 15 are at the preclinical stage. Aligned with our hypothesis, we find that as the *DKS score* of an EGFR inhibitor increases, its clinical advancement also increases (**Figure 4A**, Pearson Correlation Rho = 0.50, P = 8E-04). Furthermore, the FDA-approved EGFR inhibitors have significantly higher *DKS scores* than preclinical EGFR inhibitors (P<0.004, **Figure 4A**).

**Figure 4:**
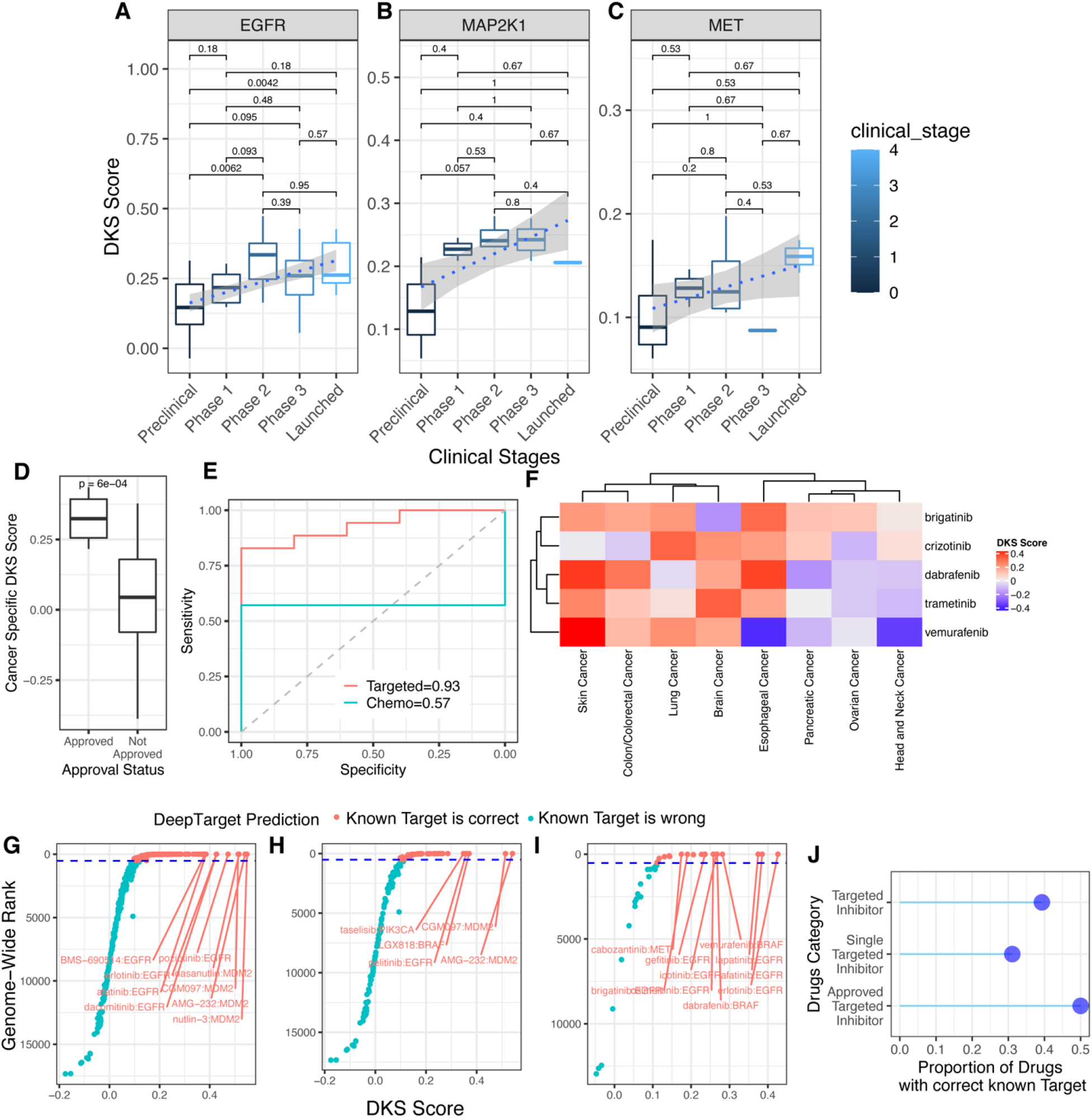
The DKS scores of the known drug targets are associated with the drug’s clinical advancement. **(A-C):**The predicted DKS score increases with clinical advancement. This is done separately for drugs targeting **(A)** EGFR, **(B)** MAP2K1 **(C)** MET. The box color intensity denoted the clinical stages (preclinical to launched). **(D-F)** A drug is more specific to its target in cancer types where it is approved. **(D)** The cancer-specific DKS score (Y-axis) of a drug’s target in approved vs non-approved cancer types and the respective stratification power is provided **(E). (F)** The DKS score of each drug in different cancer types is visualized in a heatmap. **(G-H)**. The correlation between drug response and known target CRISPR-KO for **(G)** all drugs, **(H)** for drugs with single targets, and **(I)** for FDA-approved single-target cancer inhibitors. The names of drugs whose known targets are ranked within the predicted top 10 targets by DeepTarget are specified in **(A). (J)** The proportion of drugs in each of these (A-C) categories (Y-axis) that passes the DKS score threshold (X-axis), and hence DeepTarget predictions align with their known targets.

We next tested this hypothesis for all the drugs in our screen (N=1450) and observed that, indeed, clinical advancement increases with the *DKS score* of a drug to its known target (Regression Coef = 4.32, P = 7.53E-10). This association further strengthens when we solely focus on targeted therapy drugs (Regression coefficient = 4.6, P = 5.96E-08). We next compute this relationship individually for 13 different targets with multiple drugs in various clinical phases. For 11 out of 13 targets, the clinical advancement significantly increased with their DKS scores (**Figure S6**). We illustrate these results further for the two targets with the highest number of drugs: MAP2K1 and MET, in **Figure 4B-C**, respectively. This analysis demonstrates the utility of DeepTarget to predict the likely clinical advancement of a drug.

#### The DKS score of a drug to its known target is higher in cancer types where it is approved

We next hypothesized that a drug would be more specific to its target in the cancer types where it is FDA-approved vs. the rest of the cancer types. We tested this for targeted therapy drugs (N=10) whose known targets were confirmed with DeepTarget in a pan-cancer analysis. We found that DeepTarget predicts, however, weakly statistically significant, that these drugs are more specific to their known target in their approved vs. non-approved cancer types (P=0.10, FC=1.5). A more in-depth analysis revealed that this observation holds for targeted drugs for 5 out of 6 different MOAs tested and not present for chemotherapy drugs (P=0.63). When tested for different targeted therapy MOAs separately, these observations hold for 5 out of 6 MOAs (Figure 4F). Interestingly, for targeted therapies aside from ones targeting EGFR, the drug specificity to their known target is 6.7 times higher in approved vs. non-approved cancer types (**Figure 4D**, P=2E-05, mean specificity = 0.32 vs. 0.04, respectively). This score can stratify the indications approved for a drug from all the possible cancer types (N=8) with an AUC of 0.93 (**Figure 4E**). This analysis demonstrates the potential utility of DeepTarget to predict indications likely to receive approval for a given drug of interest (**Figure 4F**).

#### *DeepTarget* predictions align with the targets currently known for about half of FDA-approved cancer drugs

We next computed DeepTarget’s DKS scores for all the target/s reported in DrugBank (its known target) for all the small molecules in the PRISM screen (N=1450, **Table S3**). This set is different from the gold-standard dataset extracted from DrugBank, as it is solely comprised of pharmacologically active drug-target pairs. Focusing on the drugs that are targeted therapy inhibitors, we observe that the known target of only 40% of drugs (N=143 out of 364) passes the DeepTarget DKS score threshold (**Figure 4F-I, Figure S7A-C**). Notably, 34 of these targets are ranked 1st in our analysis. The proportion of known targets whose predictions by DeepTarget are the same as those known is 50% and 31% for FDA-approved subsets and single-target drugs, respectively (**Figure 4J**). The distribution of the *DKS scores* for drugs in our screens (N=1450) stratified by targeted, chemo and non-cancer drugs is provided in **Figure S8**. As expected, we observed a strong enrichment in top-ranked DeepTarget predictions for known targets of targeted therapies vs. chemo and non-cancer drugs. Testing this at a pathway level identified by finding the top MOA pathway enriched in top-ranked genes yielded similar results as above (**Supplementary Notes 4, Figure S9A-C, S10A-D**). In summary, known targets of only approximately half of the FDA-approved cancer drugs align with DeepTarget predictions.

Additionally, DeepTarget predicted an alternative target for a subset of these drugs where DeepTarget predicts that their currently known target is incorrect (N=62, **Figure S7-S12**) or that they have no currently known targets (N=17, **Figure S13**), provided in **Supplementary Notes 5-6**. We provide these novel target predictions as a resource for readers to hopefully accelerate the development and repurposing of these drugs (**Table S4**).

#### Predicting drugs that target undruggable cancer oncogenes and their mutant forms

Our next goal was to predict candidate drugs that can target ‘undruggable’ oncogenes and those that can specifically target their mutant forms. To this end, we first identified oncogenes from COSMIC (N = 298) that are currently not targeted by existing drugs (N = 69). We next predicted drugs that specifically target each oncogene. We identified drugs with a DKS score for the oncogene higher than our pre-determined threshold (>0.23) and where the oncogene is ranked highest among possible targets of the drug. This yielded a few significant hits passing the DKS score threshold (**Figure 5A, S14**), where the top drug hits can target oncogenes MDM2, MITF, PPM1D, and KAT7 are shown in **Figure 5A-E** [15-18] (**Supplementary Notes 7**).

**Figure 5:**
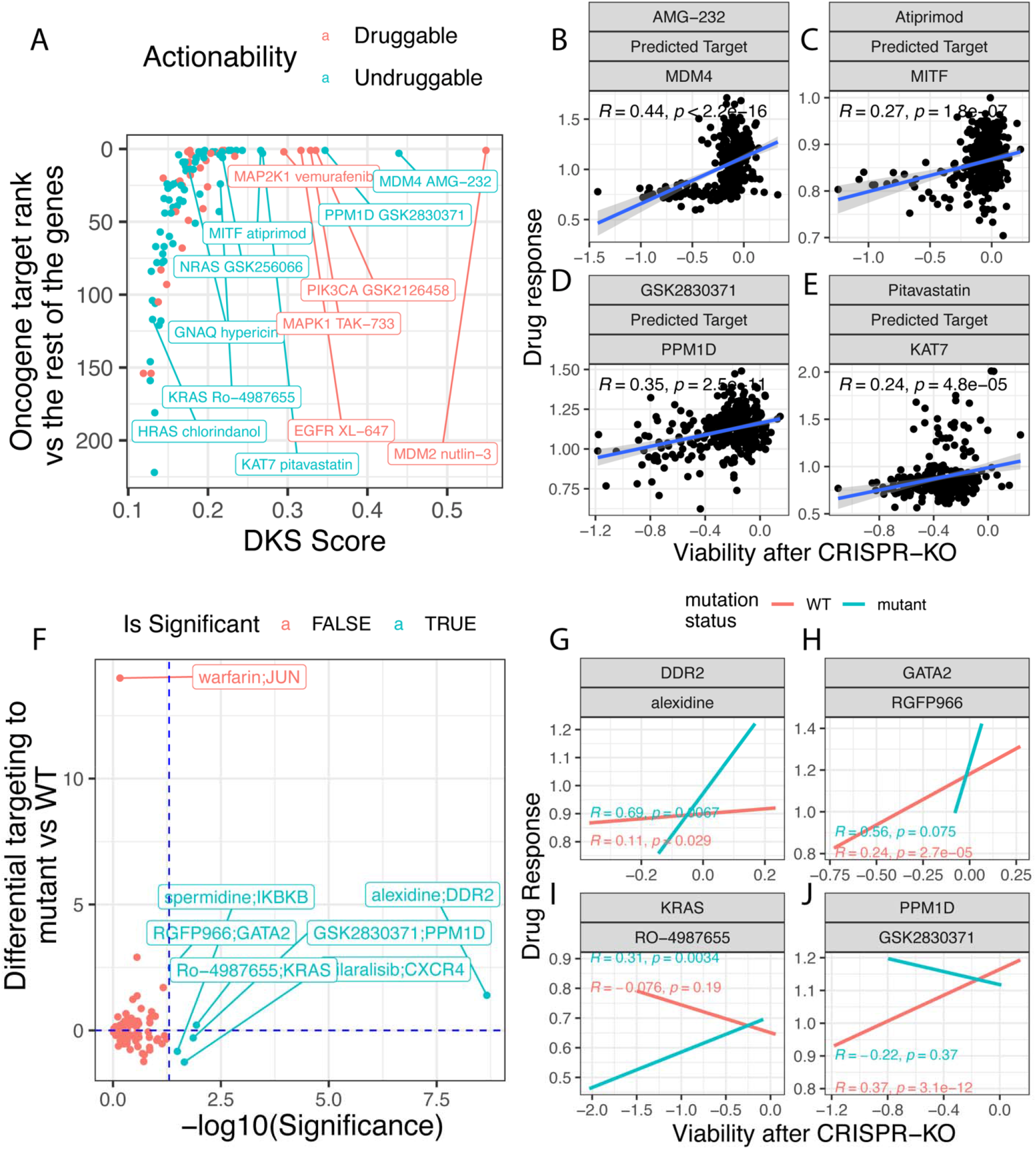
Predicting drugs targeting undruggable oncogenes and their mutant form. **(A)** The DKS scores (X-axis) of drug-oncogene pairs and the rank of the respective oncogene among all the possible targets for the drug (y-axis). The undruggable oncogenes are colored in green, and the top hits are labeled. **(B-E)** The correlation between viability after drug treatment (Y-axis) and after the KO of the predicted target (X-axis) for top drug hits. The oncogene name and the predicted drug are provided in the top strips. The strength of correlation and regression line (blue) is provided. **(F)** The mutant-specificity scores for the above drugs (Y-axis) and their respective significance (X-axis). Statistically Significant (P<0.05) hits are colored green and labeled. The vertical dashed blue line shows the statistical significance threshold. The drug above the horizontal line specifically targets the mutant form more than the WT form and vice versa. **(G-J)** Drug response to four top drug hits (Y-axis) vs. viability after knocking down the target oncogene KO (x-axis) in cell lines harboring the mutant form vs. those harboring the WT oncogene. The color legend is provided at the top, where the green and orange indicate the mutant and WT forms, respectively.

One interesting notable prediction in this category is RO4987655, a phase 1 small molecule for advanced solid tumors, is a notable hit and is predicted to target a major cancer oncogene KRAS (*DKS Score*=0.22, below but very close to TrueTarget’s decision threshold, P=0.01). This hit is interesting because, among the hits noted above, only RO4987655 is predicted to specifically target a mutant form of the target oncogene more than it is predicted to target its WT form. DeepTarget predicts that RO4987655 specifically targets the G12D mutant form of KRAS more than the KRAS WT form (**Figure 5F, 5I**, DKS Score= 0.31). Notably, the mutant form G12D KRAS is the topmost predicted target among all the genes (gene rank=1) of RO4987655. Other notable hits (**Figure 5G-H, J**) preferentially targeting the mutant oncogene form are alexidine targeting DDR2 (**Figure 5G**) and RGFP966 targeting GATA2 (**Figure 5H**).

## Discussion

Agnostic to information such as drug’s 2D/3D structures, pre/post treatment expression, adverse effects, and binding profiles, DeepTarget is based on the reasoning that the CRISPR-KO of a given drug target gene can mimic the inhibitory effects of that drug, and thus a target can be identified by identifying a gene whose CRISPR KO is a phenocopy of the drug treatment. We applied this principle in a large-scale for ∼1500 drugs and searched for their targets genome-wide by leveraging & integrating large-scale genetic and drug screens performed in cancer cell lines, markedly extending the scope and depth of the results shown in [5]. Here we comprehensively test and show the ability of DeepTarget to elucidate a drug’s: (1) primary target, (2) whether it binds more specifically to WT or mutant form of the target protein, and finally, (3) the secondary target/s that primarily mediate the response in cell-lines where the primary target is not expressed. DeepTarget is then applied to study a few fundamental standing questions in the realm of cancer drugs’ mechanism of action, as shown above.

DeepTarget has a few limitations that should be acknowledged and improved in the future. First, the results presented only refer to the drugs and small molecules whose screening data is in PRISM. However, the method itself is applicable to studying more drugs, given new pertaining screening data. Second, the RNA expression of a gene, while overall quite correlated with its protein expression levels, is, of course, known to deviate from the latter for different genes in different cancer indications. Thirdly, the expression of many genes is correlated, which may lead to background noise and various spurious results, including indirect correlations. Fourth, considering we are using viability screen with no non-cancer cells, DeepTarget currently cannot identify drug MOAs dependent on non-cancer cells and its non-viability related MOAs. Fifth, notably, the screens data is quite noisy, with significant but only moderate correlations between the results of different large-scale screens [1]. To mitigate the noise specific to CRISPR-Cas9 screens, we additionally repeated all of our analysis using RNAi screens. This reassuringly yielded overall concordant results across the study, however with lower predictive performance (**Figure S8**). We also note that our analysis does not account for a drug’s transport into the cytoplasm, specifically ABC transporters that are known to mediate drug sensitivity in cell lines [1]. Considering these different limitations and simplifications made accordingly, it is quite remarkable that DeepTarget achieves the level of prediction accuracy shown throughout this paper.

This study provides further computational confirmation for a rich list of previously known drug-target pairs. Importantly, it provides many new drug-target predictions for currently undruggable genes and for specific mutant forms. While we went to great lengths to comprehensively test the predictive ability of DeepTarget via unseen and independent datasets on an unprecedented scale, the experimental testing of specific predictions and elucidating their MOA is, of course, further deemed. It is a toll order that is out of the scope of the current investigation, but we are confident that many specific predictions outlined here for the first time will garner a broad interest and prompt such studies by others.

## Methods

### 1. Data Collection

We collected the viability screens after CRISPR-Cas9 and drug treatment from the DepMap database: https://depmap.org/portal/. The drug response screens called PRISM comprise 1450 cancer drugs and genome-wide CRISPR-Cas9 knockout viability profiles (CRISPR-KO essentiality, AVANA) of ∼17k genes, which were commonly performed across 371 cancer cell lines. The expression of these 371 cell lines was also downloaded from the DepMap portal.

### 2. The DeepTarget Pipeline

Based on the hypothesis that the CRISPR-Cas9 knockout (CRISPR-KO) of the target of a given drug can mimic the viability effect after the treatment of that drug, the viability after the target gene across multiple cancer cell lines should be similar to that viability observed after treatment with the said drug treatment. Based on this notion, the *DeepTarget* pipeline comprises three main steps: (Step 1) Predicting the primary Target of the drug, (Step 2) Predicting whether the drug binds more to the wildtype or the mutant form of the primary target, and finally, (Step 3) predicting the secondary targets of the drug that mediate the response in cell lines where the primary target is not expression.

#### 2.1 Step 1: Predicting Primary Target

In a genome-wide search, DeepTarget compares a given drug’s viability profiles across 371 cell lines to the viability profile of each gene and computes a similarity strength score (Pearson Correlation Rho). The higher the score, the higher the likelihood that the gene is the target of the drug. We termed this similarity score the Drug-KO Similarity score or DKS score. This score can range from -1 to 1, where 1 signifies a perfect correlation between viability after drug treatment and gene CRISPR-KO. The specificity of this framework is maximized across curated Platinum-standard datasets at a DKS score threshold > 0.23 (as depicted in **Figure 1**).

#### 2.2 Step 2: Predicting mutant vs. wildtype differential targeting

Based on the hypothesis that if a drug more specifically targets a mutant form of protein, in cell lines with that mutant form, the similarity score between viability after drug treatment and target CRISPR-KO (their *DKS score*) would be significantly higher than in the cell lines with WT protein. Linear regression between the viability after the drug and target KO across the cell lines with an interaction term with the mutation status of the target protein. The coefficient of the interaction term models this dependency of the DKS scored on the target mutation status and termed the ***mutant-specificity score***. A positive score denotes that the DKS score strengthens and increases in cell lines where the target is mutated and vice versa. The statistical significance of the coefficient determines if the coefficient is significantly non-zero. We repeated this analysis for specific mutation variants in the drug targets available in at least five cell lines for statistical power. This resulted in only one variant - BRAF V600E mutation. We computed drugs targeting BRAF if and the extent of differential targeting to the V600E variant.

#### 2.3 Step 3: Finding secondary targets

This step is based on the reason that a drug’s MOA, if any, would be mediated by the secondary target in cell lines with no primary target activity or expression. To this end, we first identify the set of drugs whose DKS score for the primary target indeed decreased in cell lines with decreased primary target expression. This is the case for 75% of the total drugs (N=1100). For these drugs, we identify such secondary targets genome-wide by ranking all the genes based on a *DKS score* computed in the cell lines where the primary target is not expressed. This score is called the secondary-DKS score, and the resulting genes with a secondary-DKS score>0.23 are predicted as secondary targets.

Our method assumes that the CRISPR experiment equally effectively knocks out all alleles of a gene. It also assumes that CRISPR-KO of a gene can knock out amplified or fused copies of an oncogene.

### 3. Collecting gold-standard datasets for validation of primary target prediction

As provided in detail below, we collected eight gold-standard datasets with high-confidence drug-target pairs from the following sources and preprocessed them to validate our pipeline’s prediction of the primary target. We only focused on drugs that have less than five targets in the gold-standard datasets. **(I)** The first database is curated of drug-target pairs where the target has a clinical resistance mutation extracted from COSMIC v91 (https://cancer.sanger.ac.uk/cosmic/download). We filtered targets with at least 50 unique patients with clinical resistance mutation (*N=16*) [9]. **(II)** We repeated this exercise for all such drug-target pairs with resistance mutation from oncoKB, a Precision Oncology Knowledge Base from MSKCC. (*N=28*, https://www.oncokb.org/actionableGenes#sections=Tx) [10]. **(III)** We next downloaded drug-target pairs with FDA Approval for a target mutation (FDA mutation-approval, N=86) again from oncoKB [10]. We considered pairs with all four levels of evidence in this database. **(IV)** A compilation of high-confidence drug-target pairs from ChemicalProbes.org. This is manually curated by their scientific advisory board (*N=24*) [11]. Pairs with a confidence score >3 (out of 4) are considered following database guidelines (https://www.chemicalprobes.org/information-centre#historical-compounds). This confidence score is computed by reviewing the publication and cellular and/or in vivo model systems. **(V)** A compilation of drug-target pairs with multiple independent reports as per BioGrid from https://thebiogrid.org/ (N=28) [12]. We only considered pairs with at least three independent reports. **(VI-VII**) A compilation of drugs categorized as inhibitors or antagonists where the target is categorized as pharmacologically active status as per DrugBank drug, i.e., the drug interacts directly with the target as part of its MOA. [13] This is downloaded from https://go.drugbank.com/ and yielded 90 inhibitors and 52 antagonists. **(VIII)** A list of highly selective inhibitors downloaded from compound libraries of https://www.selleckchem.com/ (SelecChem) curated based on their binding profile (N=142) [14].

### 4. Compiling gold-standard datasets for validation of mutant-specific targeting prediction

We collected two databases to validate our mutant-specific targeting prediction. **(I)** A curation of drug-target pairs where the target has a clinical resistance mutation extracted from oncoKB (https://www.oncokb.org/actionableGenes#sections=Tx). Here, due to the resistance mutation, the drugs target the mutant form more than the wild-type form [9]. We validated our method separately for both oncoKB tier I and II and yielded a significantly higher AUC for tier I. **(II)** We compiled all drugs where we have their binding profile with both target WT vs. mutant forms. To this end, we extracted binding information of all the drugs in our screen, also in https://www.selleckchem.com/. Filtering further, we focused on genes that have a specific variant in at least five cell lines in our screen to have statistical power in our comparison of WT vs. mutant. This resulted in 13 genes, where only one is targeted by a drug in our screen (BRAF). We accordingly identified drugs six drugs specifically more binding to BRAF V600E mutant form and seven that bind to both WT and mutant form comparably. Our predicted mutant-specificity score can not only stratify the two types of drugs but also correctly ranks the extent of differential binding to the V600E variant.

### 5. Predicting Ibrutinib’s secondary target

To identify Ibrutinib’s primary and secondary target, we first identified cell lines without (or low) and with BTK (Ibrutinib’s known primary target) expression. We categorized cell lines lower than log2(TPM) < 2 as cell lines with low BTK expression. We compare viability after ibrutinib treatment to all the genes in cell lines with BTK vs. low-BTK, where in the former, the DKS score for BTK is greater than DeepTarget primary target threshold (DKS score > 0.23), confirming it is a primary target. However, in cell lines with low BTK, instead, the DKS score does not pass this threshold, and EGFR is the top-ranked target.

### 6. Predicting clinical advancement

We hypothesize that a drug’s DKS for its known target, denoting its specificity towards the target, would predict its clinical advancement. We note that this hypothesis makes three assumptions: 1. This framework together considers the two types of possible errors in target information: I) entirely wrong target, II) claimed target is correct but non-specific as is often the case for kinase inhibitors. 2. Our analysis is confounded by the time period since the discovery of the drug and the start of phase I trials because a drug may not have advanced solely because it is new. We did not correct this as our screen does not comprise recently discovered drugs, and most of the drugs in our screen have been in trials or a preclinical stage for long enough to mitigate this effect. 3. We consider different generations of drugs equivalently and do not consider that a latter generation may have a predecessor generation drug with effectiveness. In the case of drugs targeting EGFR, we consider all drugs targeting EGFR, whether they target the specific WT or mutant form and different generations equally.

### 7. Predicting cancer types that would likely get approved for a drug

We next hypothesize that a drug would be more specific to its target in cancer types where it is approved. We note that this hypothesis assumes the following: (1) We consider a drug targeting the mutant or WT form preferentially equally. (2) We also assume that the frequency of the mutation among cell lines is a good representation of the frequency of the mutation among patients.

## Data Availability

Processed PRISM drug screen, expression, mutation, copy number, CRISPR-Cas9, and shRNA pooled genetic screen data were derived from DepMap v22Q4 and can be found here (https://depmap.org/portal/).

## Code Availability

We have provided the scripts to reproduce each Figure in the main text and supplementary figures in a GitHub repository, which can be accessed here: https://github.com/ruppinlab/drug.OnTarget

## Conflict of Interest

Authors Eytan Ruppin (E.R) and Sanju Sinha are the inventors on a pending provisional patent application related to the DeepTarget algorithm. E.R. is a co-founder of Medaware, Metabomed, and Pangea Therapeutics (divested from the latter). E.R. is a non-paid scientific consultant to Pangea Therapeutics, a company developing a precision oncology SL-based multi-omics approach. The rest of the authors declare no conflict of interest.

## Acknowledgments

We thank the National Cancer Institute for providing financial and infrastructural support. This research was partly supported by the Intramural Research Program of the National Institutes of Health, NCI.

