## Supplementary Figures and Text for "Deep characterization of cancer drugs mechanism of action by integrating large-scale genetic and drug screens"

### Graphical Abstract

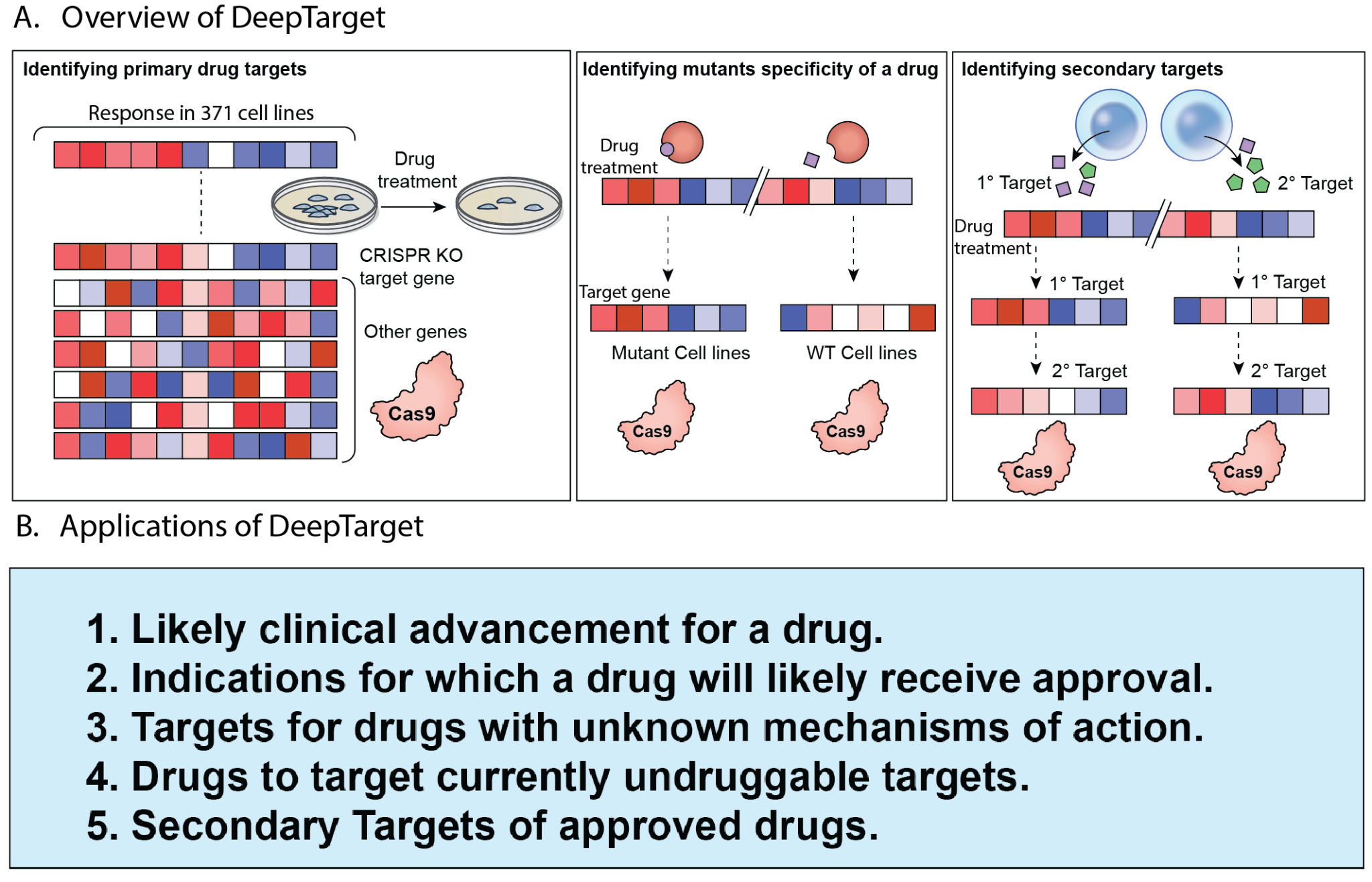

**Supplementary Notes 1: Curating Gold-Standard database for primary target prediction testing.**

A. *Curation process of seven gold standard datasets.*

The following sources are used for gold standard drug-target pairs datasets (**Table S1**): 1. Clinical resistance mutation in target from COSMIC (COSMIC resistance, N=16), or 2. oncoKB (oncoKB resistance, N=28), 3. FDA Approval for a target mutation (FDA mutation-approval, N=86), 4. High confidence as per the scientific advisory board of ChemicalProbes.org (SAB, N=24), 5. Multiple independent reports as per BioGrid (Biogrid Highly Cited, N=28), 6. pharmacologically active status as per DrugBank drug, i.e., the drug interacts directly with the target as part of its mechanism of action and are inhibitors (DrugBank Active Inhibitors, N=90) or 7. are antagonists (DrugBank Active Antagonists, N=52), 8. Highly selective inhibitors based on their binding profile (SelecChem selective inhibitors, N=142) [14].

We next filtered pairs all the pairs that had less than ten targets. We next put a qualitative filter including the following to remove low-quality data from these datasets. The pairs from BioGrid were next filtered for at least five independent citations. The pairs from COSMIC must be supported by data from at least 50 patients. The pairs from SAB are filtered for at least an average rating from the board of three. We note that both levels R1 and R2 are considered in the oncoKB resistance dataset, where the AUC observed for pairs in the qualitatively superior and more robust R1 group is higher than for pairs in R2. We also note that all four levels of clinical evidence available are considered from the oncoKB-approved dataset, where the AUC observed for groups are in order of their reliability.

B. *Subsetting drug-target pairs with high qualitative scores*

Subsettting premium quality pairs in the last three datasets noted above (*SAB*, *oncoKB resistance*, FDA mutation-approval) by utilizing a SAB average score threshold of four, subsetting to oncoKB pairs with resistance mutation information that are of clinical standard of care (R1 group only), FDA mutation-approval pairs that are either FDA approval or standard of care. Subsetting the drug-target pairs by these three criteria yields an average AUC of 0.96.

**Supplementary Notes 2: Stratification Power by different drug target Classes and restricting the search during target identification**

A. *Stratification power by target class and pathway*

We first note that the stratification power of DeepTarget to identify gold-standard drug-target labels from negative control (AUC) for different drug categories defined by their protein type is 0.82 for kinase inhibitors, 0.61 for ion channel inhibitors, 0.60 for ion channel inhibitors, 0.61 for nuclear receptors, 0.55 for proteases and 0.65 for metabolic enzymes. Computing this stratification power across individual pathways, we found that the top pathways enriched in the targets ranked by DeepTarget prediction power are PI3K/AKT Signaling (FDR<1E-14), growth factor receptors (FDR<1E-14), FGFR Signaling (<2E-13) and RAF/MAP kinase pathway (<7E-13).

B. *Restricting target space to genes likely to bind to drug*

In our analysis to mitigate the prediction of highly ranked genes with a similar KO profile (downstream or upstream of the target), we performed the target identification process for drugs with gold-standard target information but instead of using a genome-wide search space, we limited the target search space to the protein class the drug is designed towards and would thus likely bind. Here, false positives are hits ranked higher than the reported gold-standard target. While restricting to the Kinase space yielded no change in AUC, the false positive rate decreased from 13% to 0.003% of total genes. Repeating this analysis for major drug target classes yielded no decrease in AUC for each class, GPCR (AUC=0.56), nuclear receptors (0.61), ion channels (0.60), and metabolic enzymes (0.65); however, it significantly decreases the false positive rate in each class from an average 22% to an average of 0.001%.

**Supplementary Notes 3: *Identifying known secondary targets of drugs***

Another notable example is Tamibarotene, an approved drug in Japan against acute promyelocytic leukemia and investigated as a possible treatment for Alzheimer's disease, multiple myeloma, and Crohn's disease. *DeepTargets* uncovered its previously known targets: retinoic acid receptor, alpha (RARA, Rank=264), and beta (RARB, Rank=284) as the primary and secondary targets, respectively (**Figure S13**). An analysis focusing on all cell lines could not uncover the secondary target retinoic acid receptor, beta (Rank=10,504). Another such example is AZD5363 and fludarabine, with AKT1 & AKT3 (Rank=12 & 513) and RRM1 & RRM3 (Rank=70 & 239), as the primary and secondary targets that can only be identified using the combined two-step *DeepTargets* analysis.

**Supplementary Notes 4: MOA identification at Pathway Level**

#### Testing at a pathway level, we computed and tested whether the drug’s mechanism of action pathway is enriched at the top among the predicted ranked target list from DeepTarget. Focusing on targeted therapy inhibitors, including multiple targets, we found that for 78 out of 364 drugs (proportion= 21%), the known mechanism of action passes DeepTarget’s threshold (**Supplementary Figure 3A, 3D**). This analysis for all drugs focusing on single target drugs is provided in **Supplementary Figure 3B** and only FDA-approved ones in **Supplementary Figure 3C**. This proportion is 34% and 19.5% for FDA-approved subsets and single-target drugs (**Supplementary Figure 3D**).

**Supplementary Notes 5: Predicting the correct targets for drugs with discordantly/incorrectly annotated targets**

Focusing on drugs whose publicly annotated targets are predicted as wrong by DeepTarget (N=221) or with no currently annotated targets, we computed their best-predicted targets by *DeepTarget.* The *DKS scores* for the annotated and the best-predicted targets are shown in **Figure 3A-B (Figure S3B** shows this for all 1450 drugs). For each drug, we quantified the improvement of the *DKS score* for our best-predicted target vs. the one currently annotated, termed its *Improvement score* (**Figure 3C**). Among these drugs, there are 75 drugs for which our pipeline can predict an alternative novel target with high confidence (Correlation strength FDR P<0.1 & improvement score >0.2, highlighted in blue in **Figure 3A**). The distribution of the clinical trial stages of these drugs is provided in **Figure 3D**. The pathway annotation of the target that is predicted to be incorrect and the newly predicted one is provided in **Figure 3F.** The respective overall pathways enrichment is shown in **Figure S10A-B**. We also repeated a similar analysis at the pathway level instead of the gene level and observed a comparable landscape (**Figure S3C-E**).

One of our top hits is strophanthidin, whose annotated target is *ATP1A1*, a cation transport ATPases. The correlation between strophanthidin response and ATP1A1 CRISPR-KO is 0.09 (*DKS Score*, **Figure 3E**). In contrast, our pipeline predicts strophanthidin is a proteasome inhibitor targeting PSMD5 (*DKS Score*=0.51, **Figure 3F**). We provide the novel targets for these 75 drugs and their *DKS scores* as a resource in **Table S3**. Among the FDA-approved drugs, our pipeline predicts that Ribociclib, an FDA-approved drug for breast cancer treatment and annotated as a CDK4 inhibitor, instead targets the biosynthesis enzyme of phosphatidylinositol, a downstream messenger of many G protein-coupled receptors and tyrosine kinases regulating cell growth (**Figure S10C,** *DKS Score* of CDK4=-0.004 vs. that of CDIPT=0.22, P=1.5E-05). We also predict that decitabine, a DNA hypomethylation agent considered to act via DNMT1 inhibition, instead inhibits CFAP20 (**Figure S10D,** *DKS Score* DNMT1= -0.04, CFAP20=0.27, P=3.6E-07).

**Supplementary Notes 6:** Predicting the correct targets for cancer drugs with no currently known targets

We next turned to predicting the targets of cancer drugs with no currently reported targets (N=58) included in the screens. *DeepTarget* identifies a statistically significant target for 17 drugs (FDR P<0.05), whose pathway annotations are provided in **Figure S11A**. Among these, one of the top-ranked drugs is BVD-523, a cancer investigational drug in phases 1-2 with no target, which is predicted to inhibit Cyclin Dependent Kinase 2 Interacting Protein (CINP, DKS=0.29), which is also known to serve as cell cycle checkpoint regulator (**Figure S11B**). Another notable hit is Pibenzimol, a fluorescent molecule known to bind to double-stranded DNA and in clinical phase 2 due to complete remission in a single pancreatic patient and low toxicity [15]. *DeepTarget* predicts that it inhibits the E2 ubiquitin-conjugating enzyme family (**Figure S11C,** UBE2Q1, *DKS Score*= 0.27, P=1.5E-07). Thirdly, *DeepTargets* predicts that Iobenguane, a radiopharmaceutical agent with no protein target and recently approved for rare tumors [16], inhibits Contactin 3, an immunoglobulin expressed exclusively in the nervous system that regulates neurite growth (**Figure S11D,** *DKS Score*= 0.25, P=1.6E-04). A fourth hit is Temoporfin, a currently EU-approved photosensitizer, which is predicted to inhibit a nucleolar phosphoprotein *NCL* involved in the synthesis and maturation of ribosomes (**Figure S11E**, *DKS Score*=0.23, P=1.2E-05).

Supp Notes 7: Drugs predicted to target undruggable oncogenes

Our interesting top hits include BMS-707035, a phase 2 drug being tested in HIV patients that is predicted to target MYCN, a central driver of neuroblastoma (*DKS Score*=0.21, Rank=2). Another notable hit is AMG-232, a known inhibitor of MDM2, which is also predicted to target MDM4 (**Figure 5B,** *DKS Score*=0.44, gene rank=3). We also predict that *Atiprimod* targets and inhibits *MITF*, a transcription factor involved in melanoma [19] (*DKS Score*=0.27, gene rank=3, **Figure 5C**). Our next notable hit is the small molecule GSK2830371, which is predicted to target PPM1D, a negative regulator of p53. GSK2830371 has been previously reported to demonstrate anti-tumor efficacy in cell lines overexpressing PPM1D by inducing G2/M arrest and apoptosis (**Figure 5C**) [17]. Another hit is Pitavastatin, an FDA-approved statin, which is predicted to inhibit KAT7, an undruggable oncogene involved in clear cell renal cell carcinoma (*DKS Score*=0.24, gene rank=4, **Figure 5D**).

### Supplementary Figures

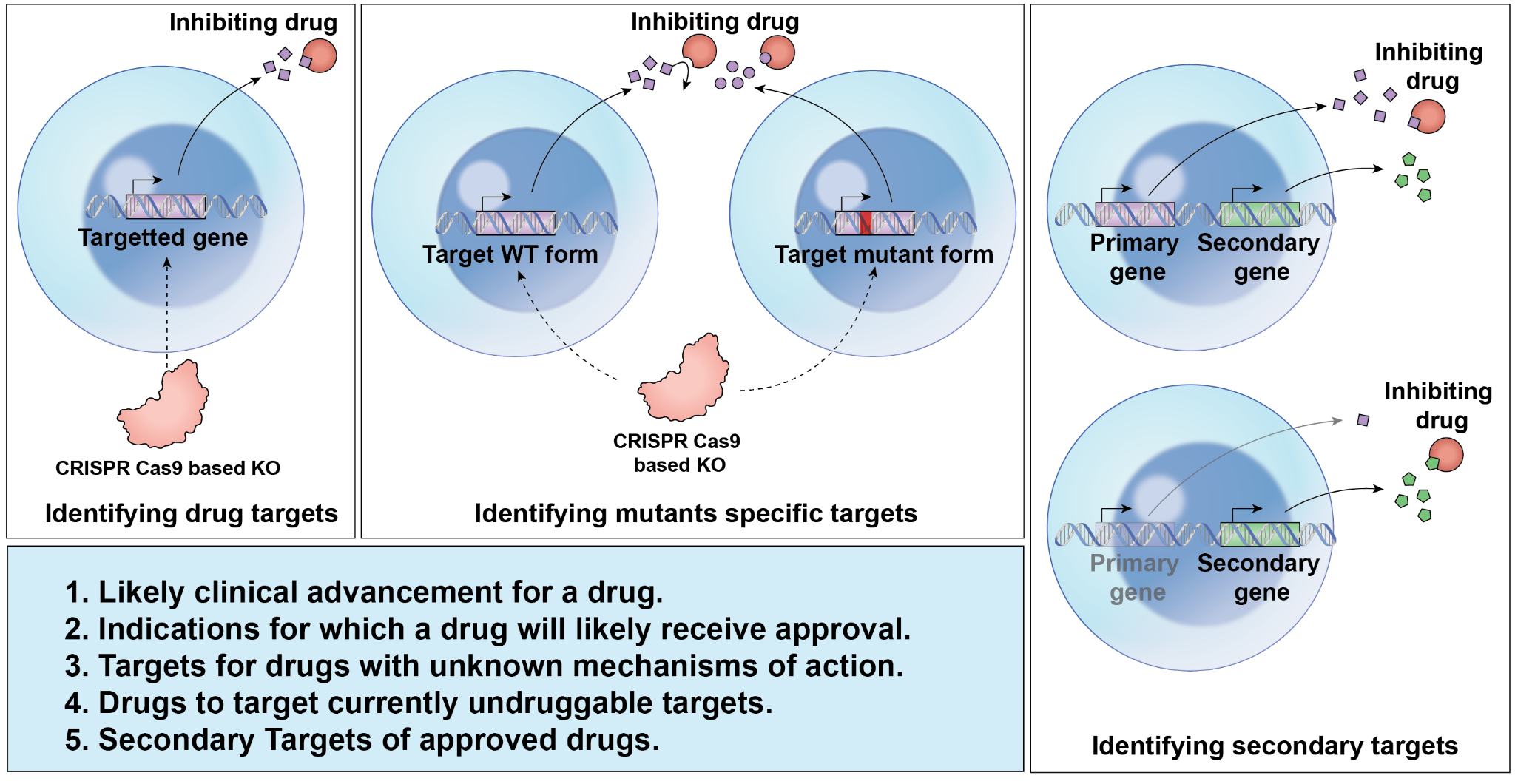

**Figure S1: The principle of DeepTarget.** Our pipeline is based on the principle that the CRISPR-Cas9 knockout of a gene encoding the protein target of a given drug can mimic the inhibitory effects of that drug, and thus a target can be identified by identifying a CRISPR knockout phenocopied by the drug. A list of DeepTarget applications is provided at the bottom.

| 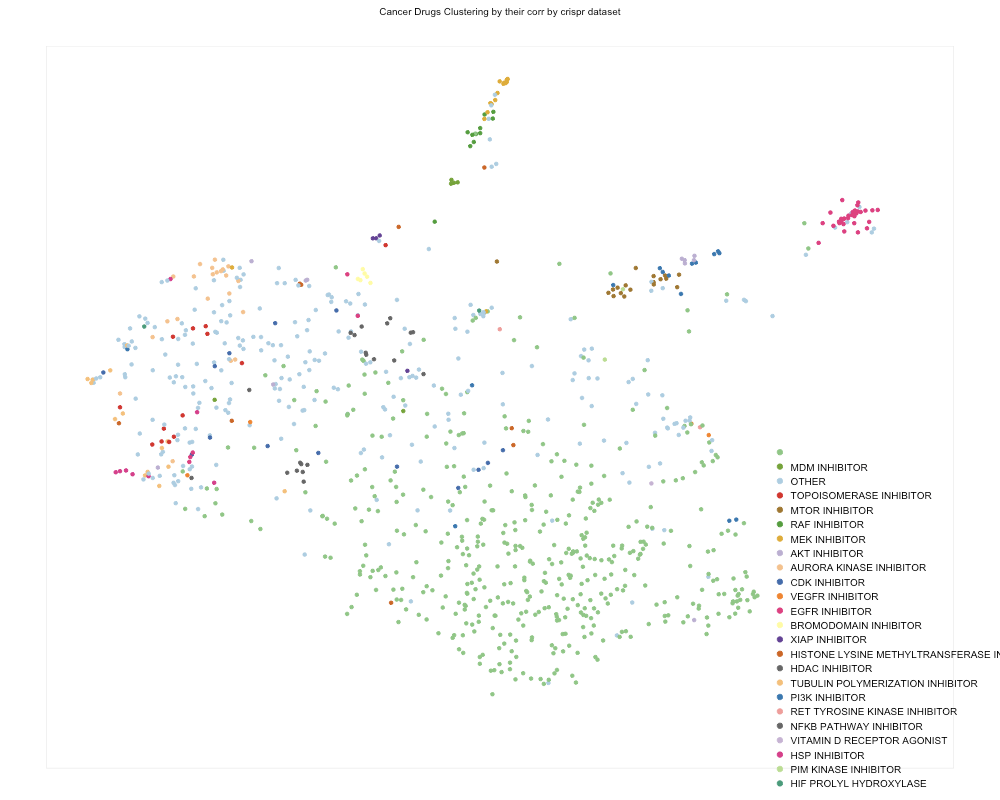 | 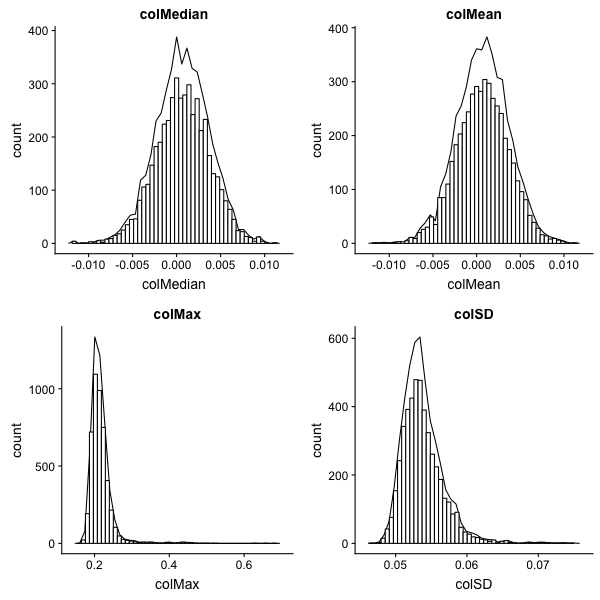 |
| --- | --- |
| 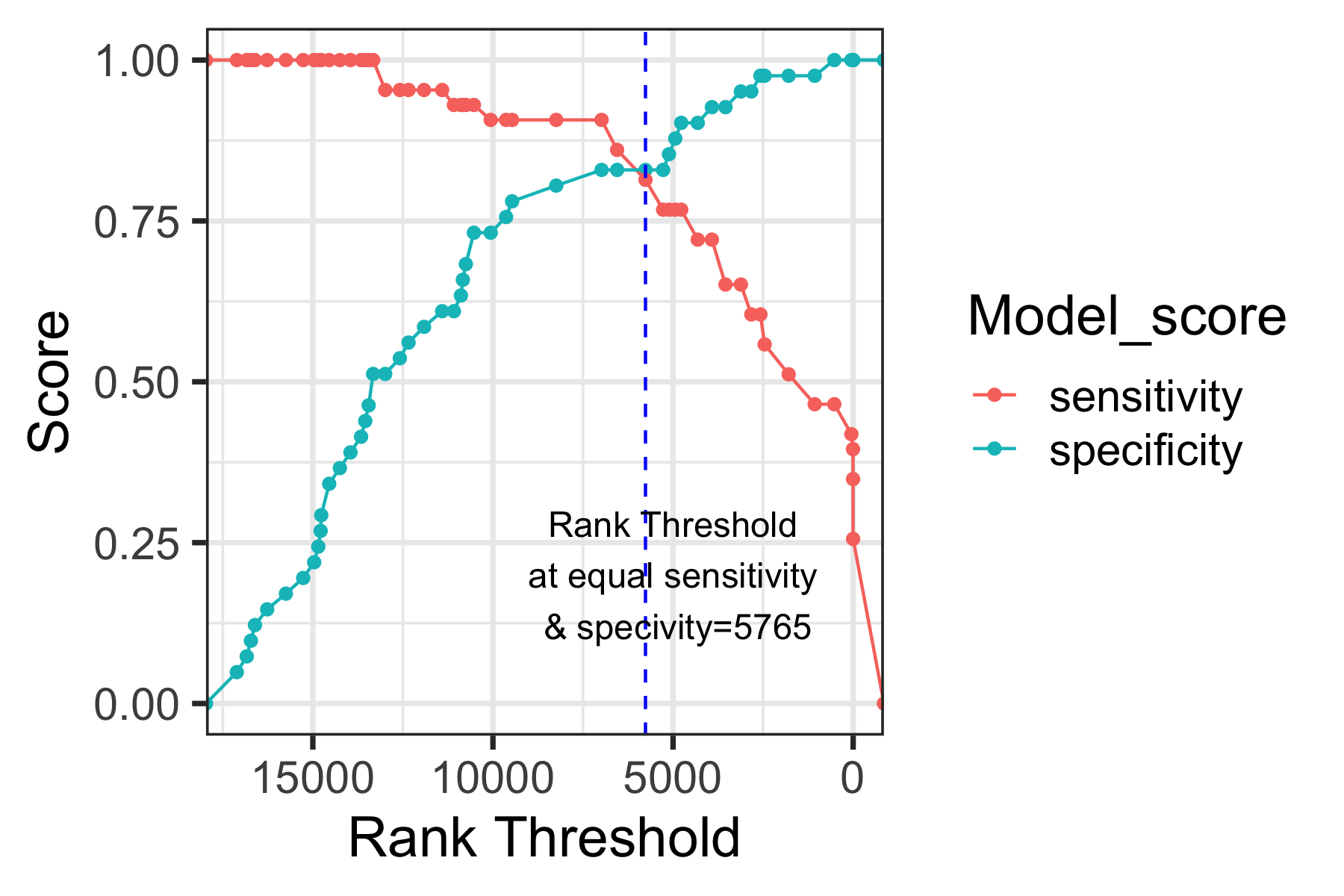 |  |

**Figure S2**: The UMAP representation of the drug-KO similarity index for all the drugs where each point represents a drug colored by their known mechanism of action. The color legend is provided at the bottom right. (c) Here, Sensitivity means the total proportion of actual targets that are recovered, and Specificity means the proportion of genes that are not a target identified as not a target.

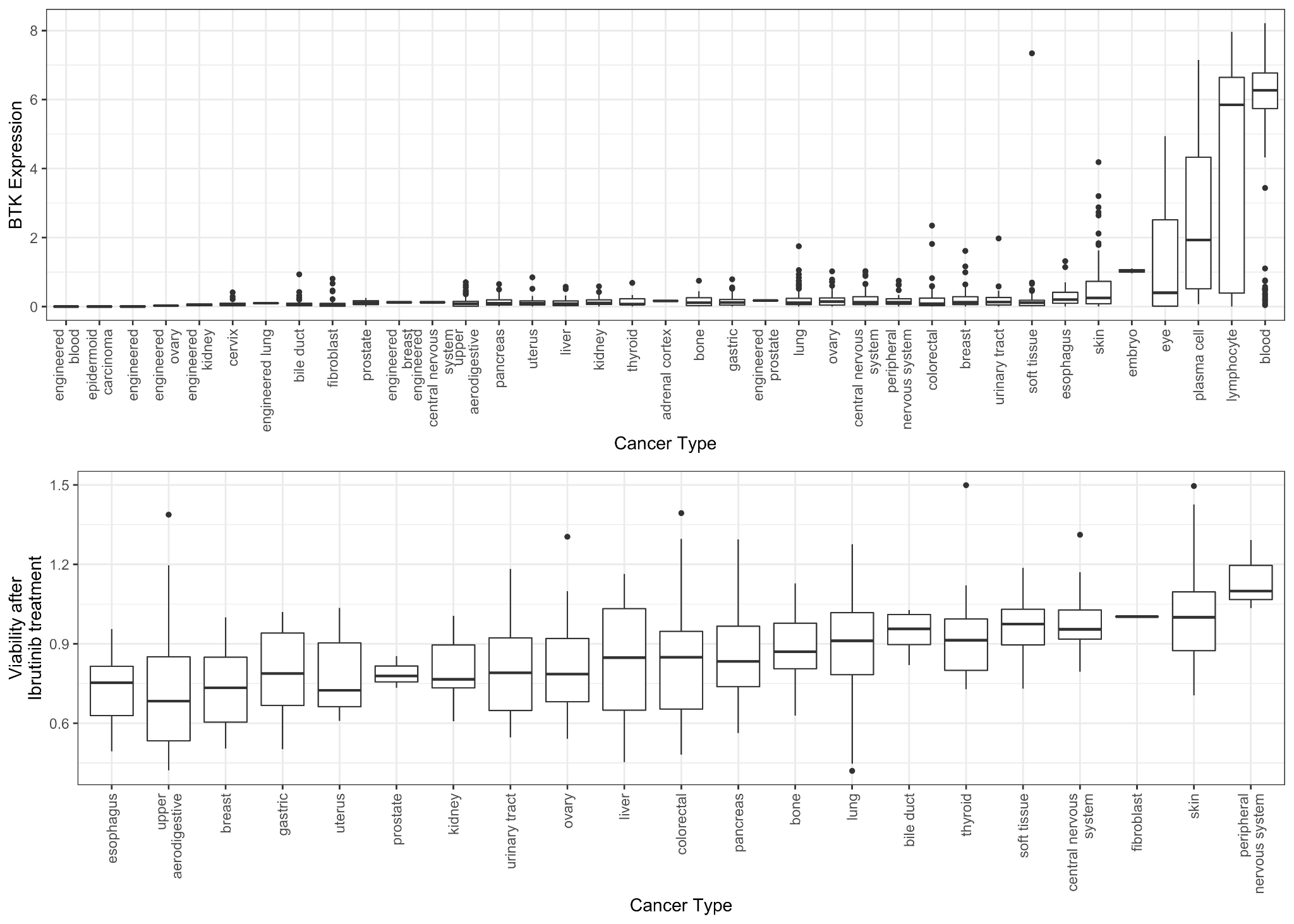
**Figure S3: Expression of BTK gene in cell lines from different cancer types.**

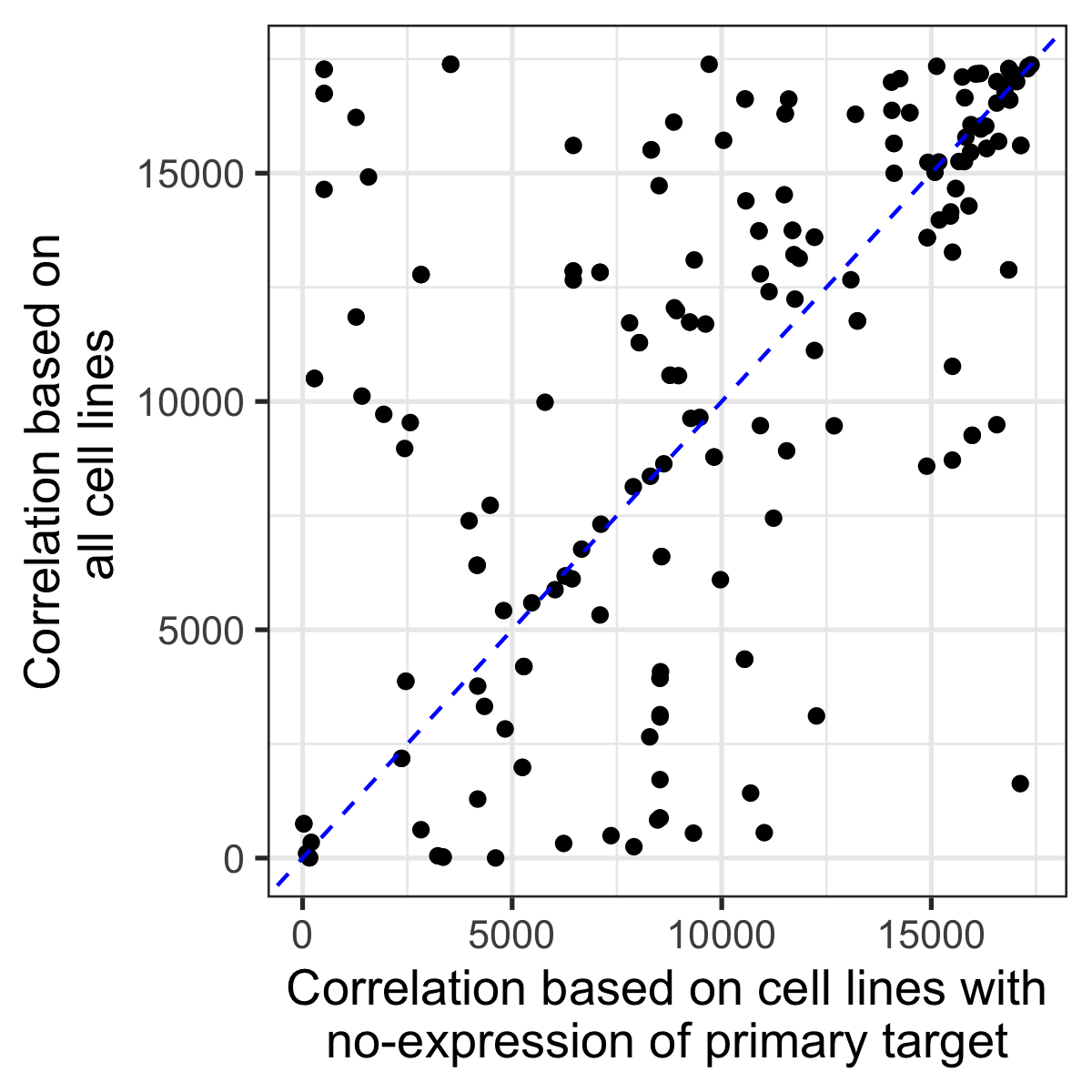

**Figure S4:** Predicted score of secondary targets considering all cell lines vs cell lines with no expression of the primary target for drugs whose primary target is corrected as predicted by our pipeline.

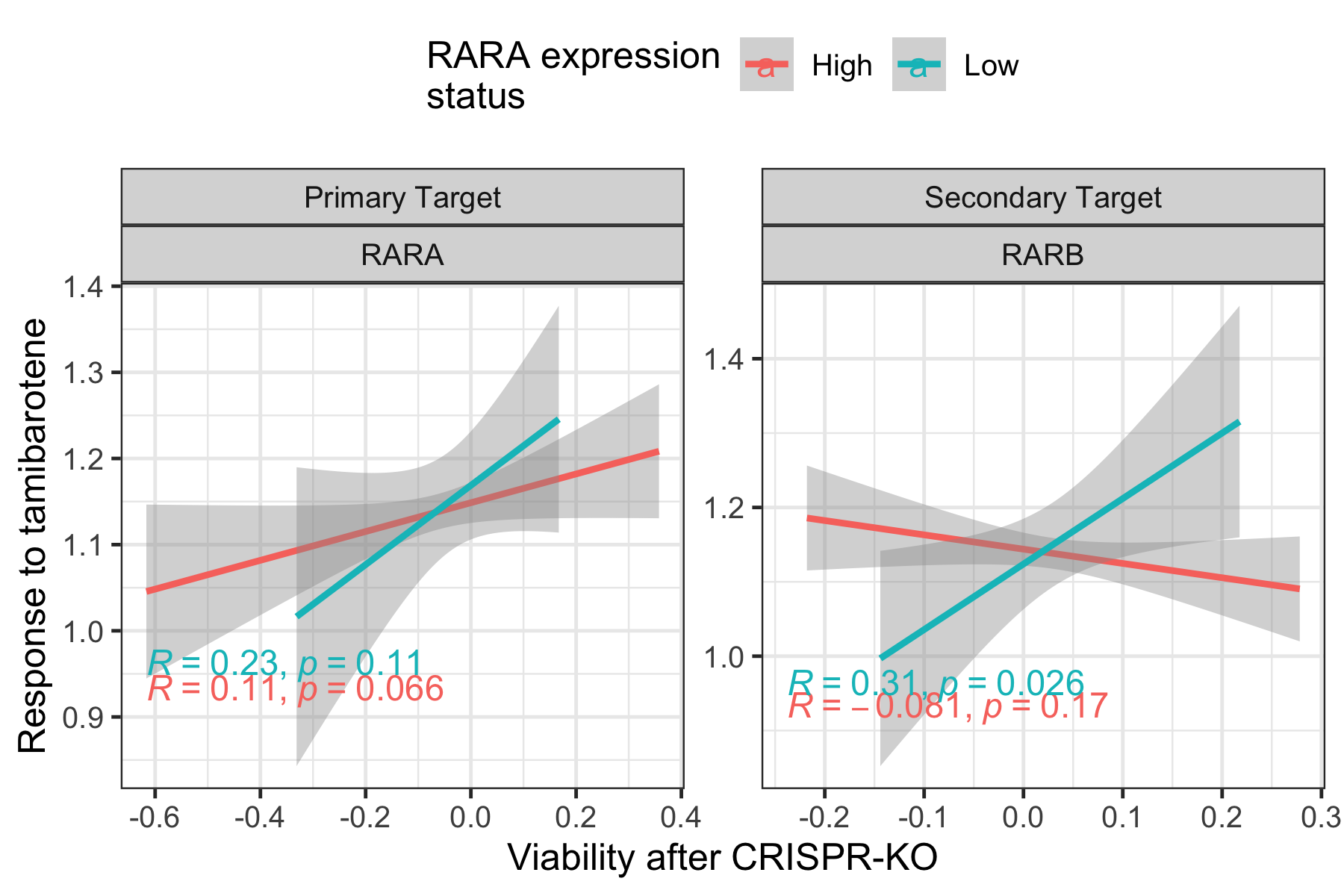

**Figure S5: The response to Tamibarotene vs viability after CRISPR KO of RARA and RARB in cell lines with RARA high vs low expression cell lines.** Tamibarotene is an approved drug in Japan against acute promyelocytic leukemia and is investigated as a possible treatment for Alzheimer's disease, multiple myeloma, and Crohn's disease. Our pipeline uncovered both previously known targets: retinoic acid receptor, alpha (RARA, Rank=264), and beta (RARB, Rank=284) as a primary and secondary targets, respectively. An analysis focusing on all cell lines was not able to uncover the secondary target retinoic acid receptor, beta (Rank=10,504).

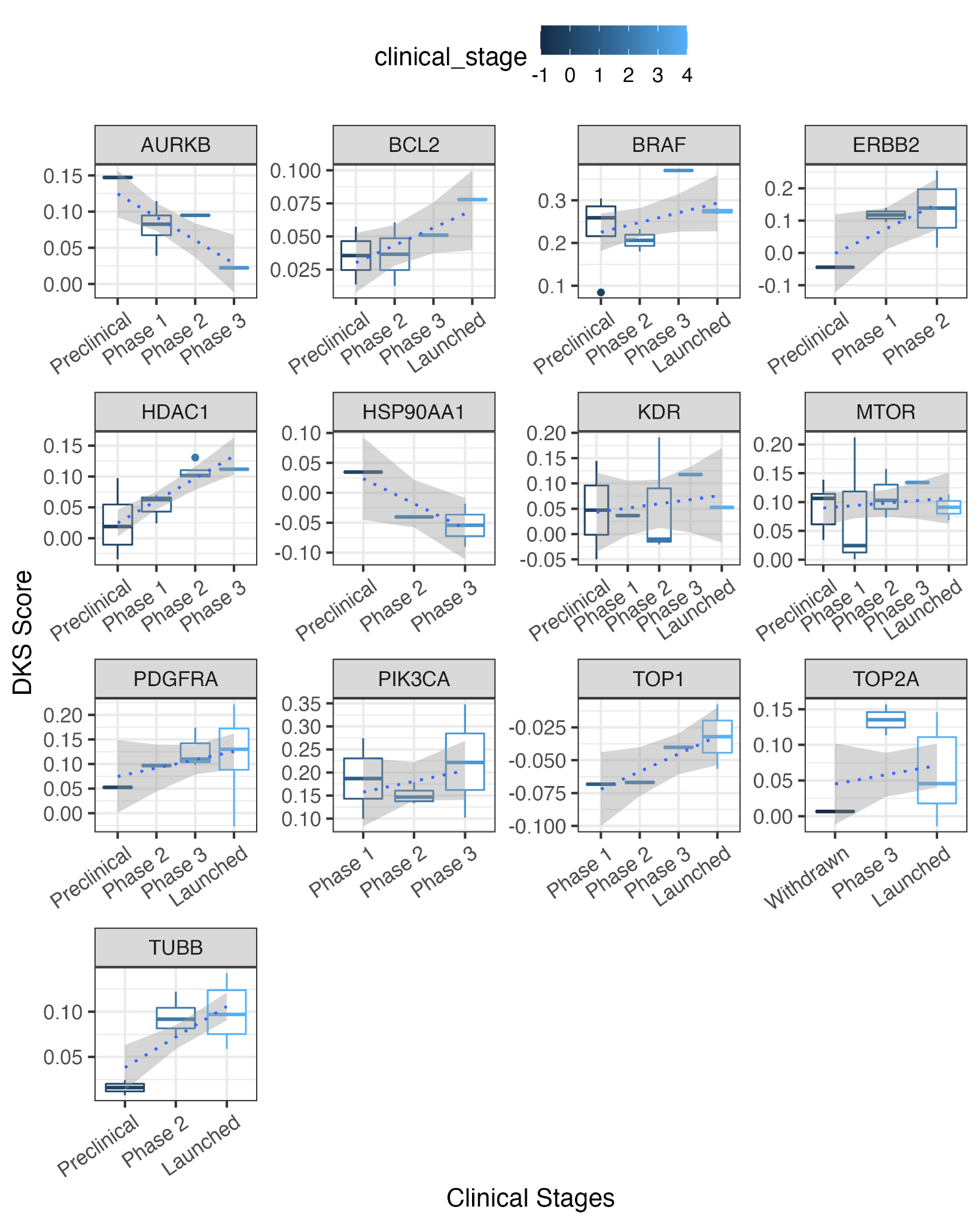

**Figure S6:** Predicting the clinical advancement of an addition 13 top targets where at least three clinical phase information is available. *Y-axis represents the DKS score and the x-axis represents the Clinical stages. The legend colors are in increasing order of clinical stages (preclinical to launched).*

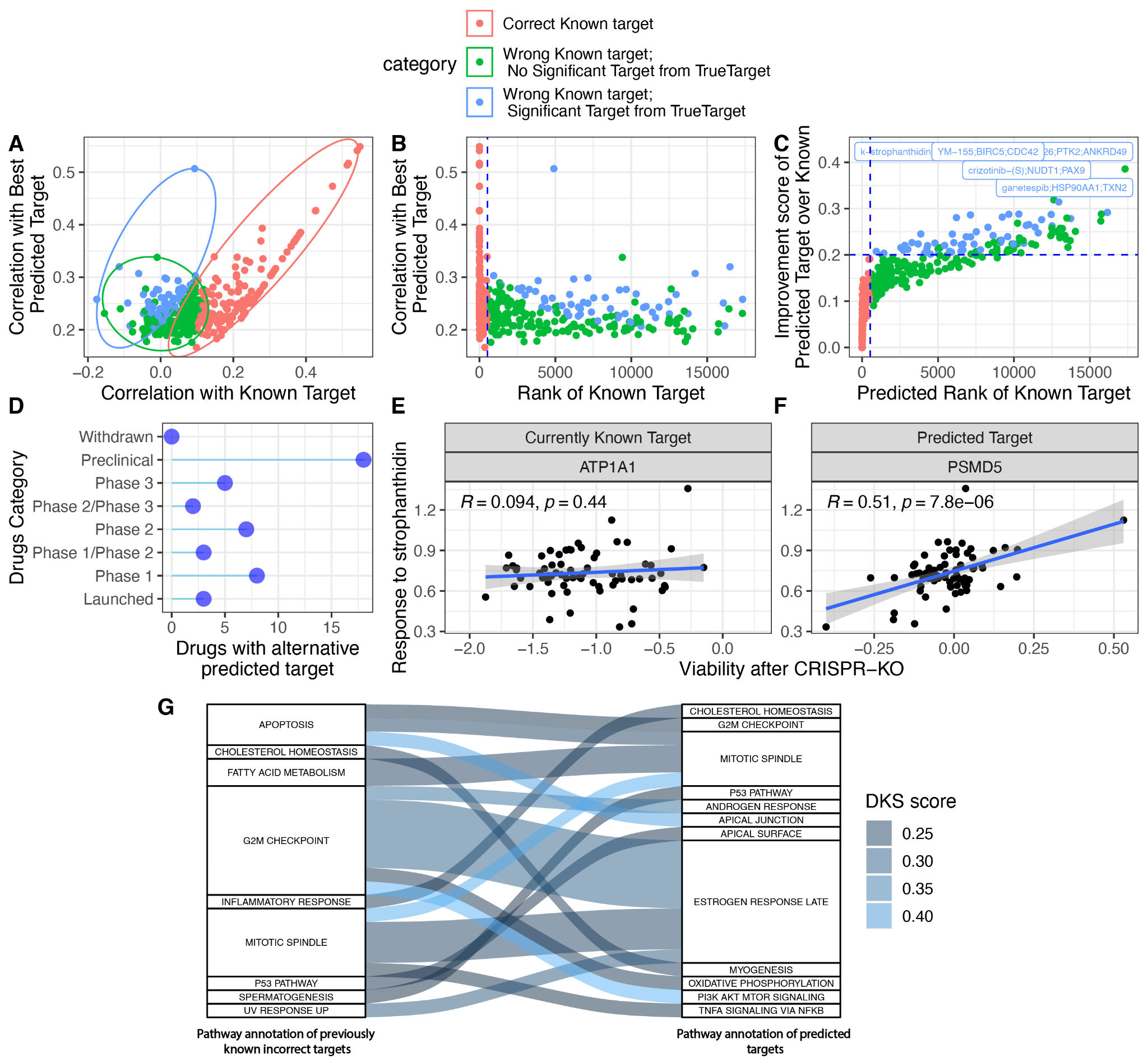

***Figure S7: Predicting new targets for drugs with targets that are predicted to be wrongly annotated or with no known targets. (A)*** *Correlation between drug response and CRISPR-KO of the currently annotated target (x-axis) vs best-predicted target by TreuTarget. The three colors represent the three categories of drugs: 1. The known target is predicted to be corrected by our pipeline (red, Predicted rank<523), 2. The known target is predicted to be incorrect (predicted rank>523) and our pipeline cannot identify any new significant target (FDR P<0.1, green), and the most important group, where 3. The currently annotated target is predicted to be incorrect (predicted rank>523) and our pipeline predicts a new target, with a statistically significant correlation with drug response (FDR P<0.1) with an improvement score>0.2 (blue).* ***(B)*** *The distribution of DKS scores of the best-predicted target (Y-axis) vs the predicted rank of the currently annotated target (X-axis).* ***(C)*** *The predicted rank of the currently annotated target (X-axis) vs the respective improvement score.* ***(D)*** *The number of drugs that are predicted to have a significantly improved target, stratified by clinical stage.* There are 18 preclinical, 11 phase 1, 9 phase 2, 5 phase 3, and 2 FDA-approved drugs. ***(E)*** *The correlation between viability after drug treatment (Y-axis) and after the KO of the currently annotated (ATP1A1, left panel) and DeepTarget predicted new target (PSMD5, right panel) across all cell lines for strophanthidin. The strength of correlation and regression line (blue) is provided.* ***(F)*** *The mapping between the pathway annotation of currently annotated targets (left) and newly predicted ones (right) is provided. The intensity of the connection represents the improvement score and the width represents the number of drugs with such a transition. The color legend is provided in right.*

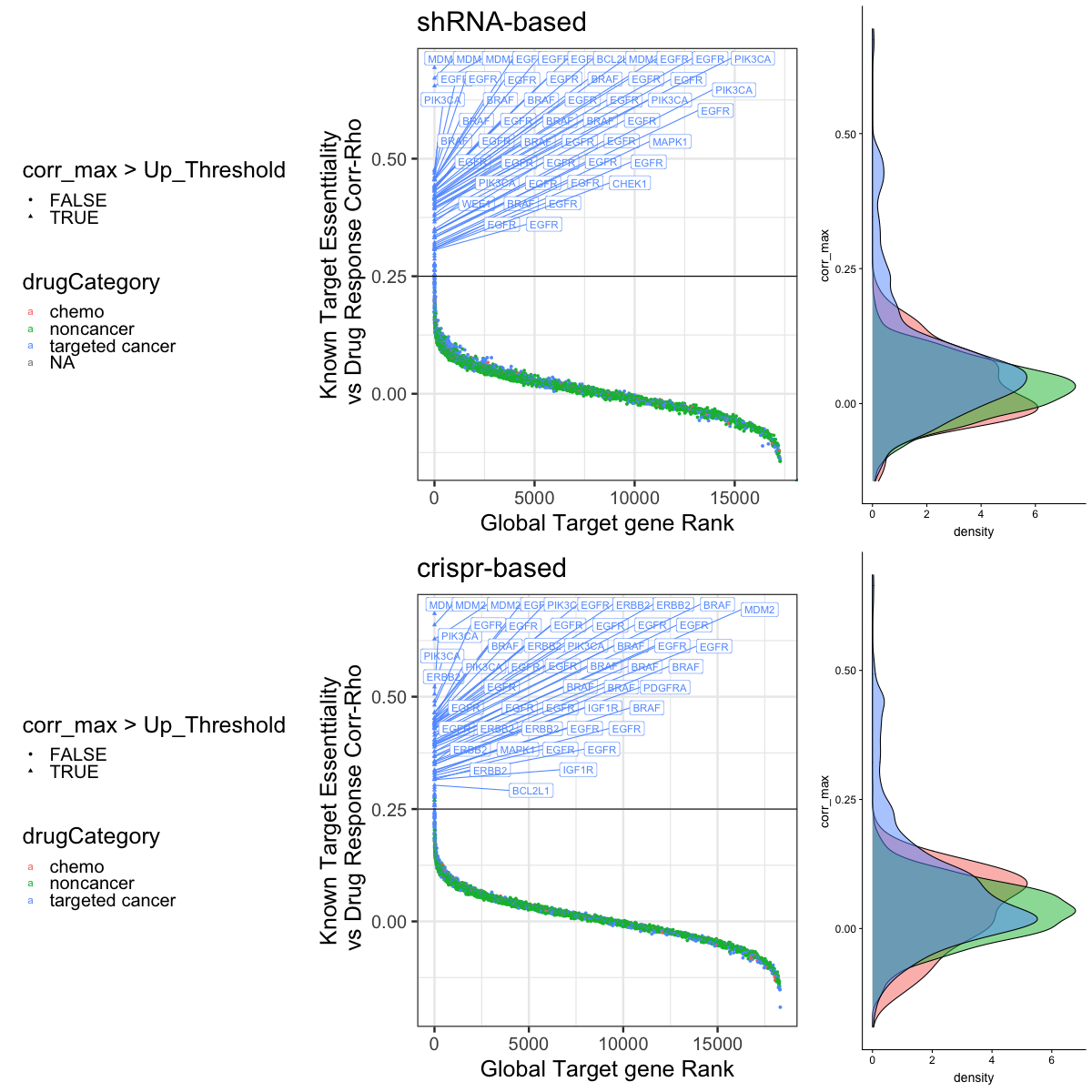

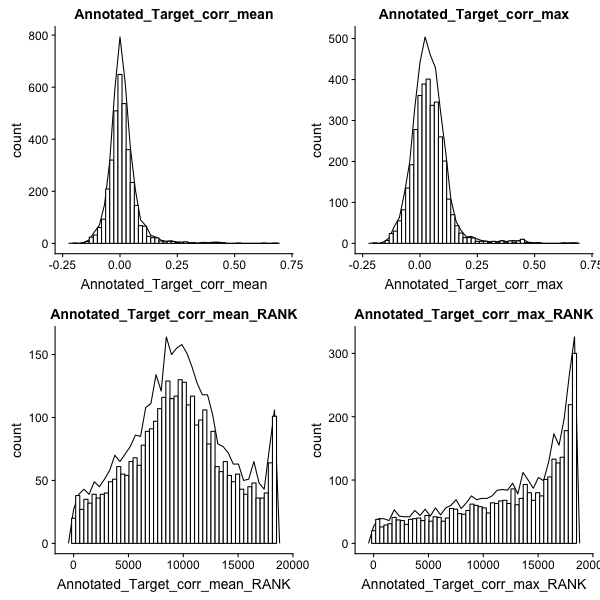

**Figure S8: The correlation predicted by our pipeline** Drug response and their target CRISPR-KO correlation strength in our screens (N=1618) stratified by targeted, cancer, and non-cancer.

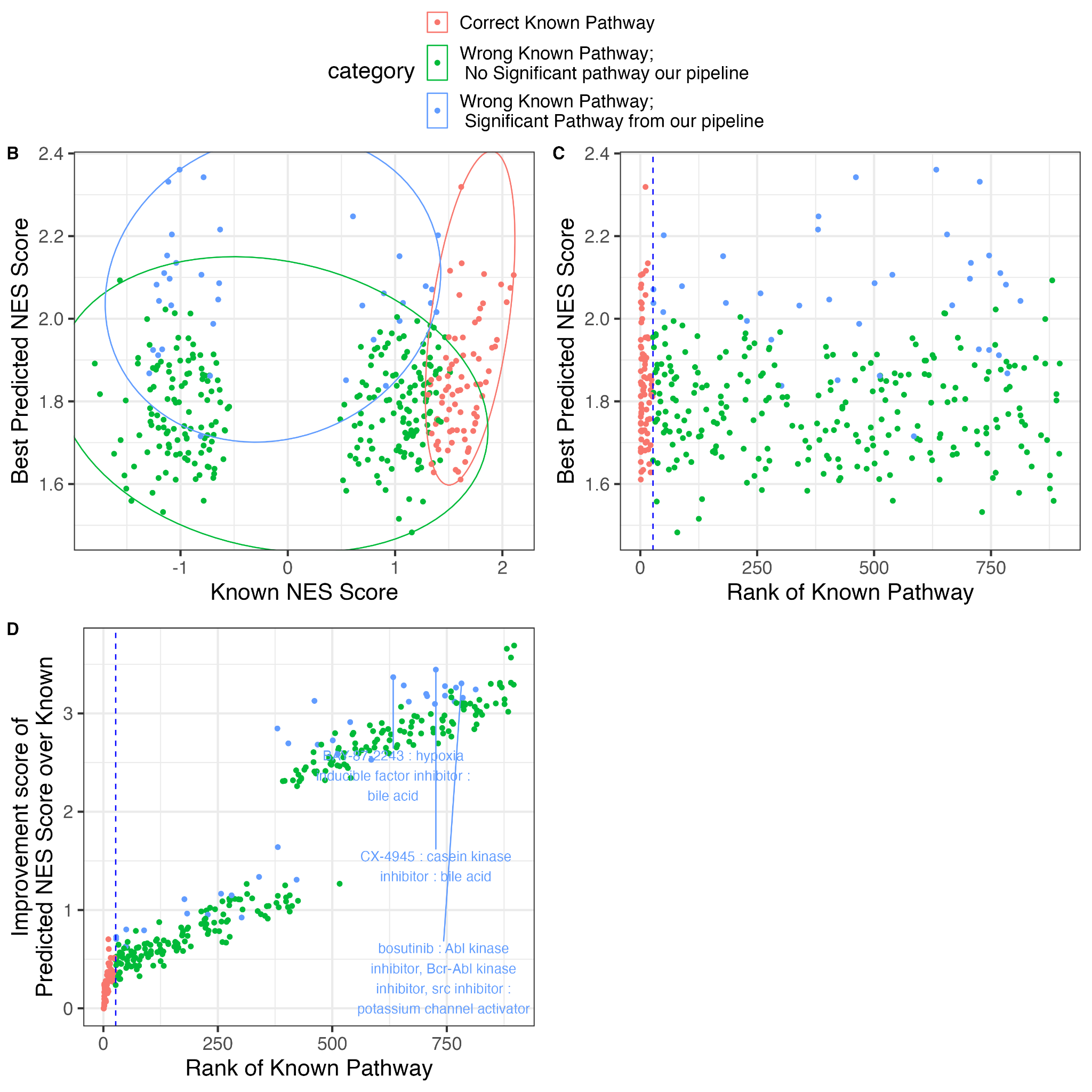

**Figure S9: *Predicting pathway for drugs with wrong known pathway or no known pathway. (A)*** *Correlation between NES Score of the known mechanism of action (x-axis) vs best-predicted mechanism of action from our pipeline. The three colors represent the three categories of drugs: 1. known pathway is predicted to be corrected by DeepTarget (red, DeepTarget pathway threshold: rank < 27), 2. known target is incorrect ( DeepTarget pathway threshold: rank < 27), and our pipeline cannot identify any new significant pathway (FDR P > 0.2, green), and the most important group where 3. the known pathway is incorrect (DeepTarget pathway threshold: rank < 27) and our pipeline can predict a statistically significant target (FDR P<0.2) with an improvement score>0.1 (blue).* ***(B)*** *NES Score of the best-predicted pathway from our pipeline (Y-axis) and the predicted rank of the known pathway from our pipeline (X-axis).* ***(C)*** *The predicted rank of the known target from our pipeline (X-axis) vs the improvement score i.e.* the difference of their known vs best hit correlation strength with the response.

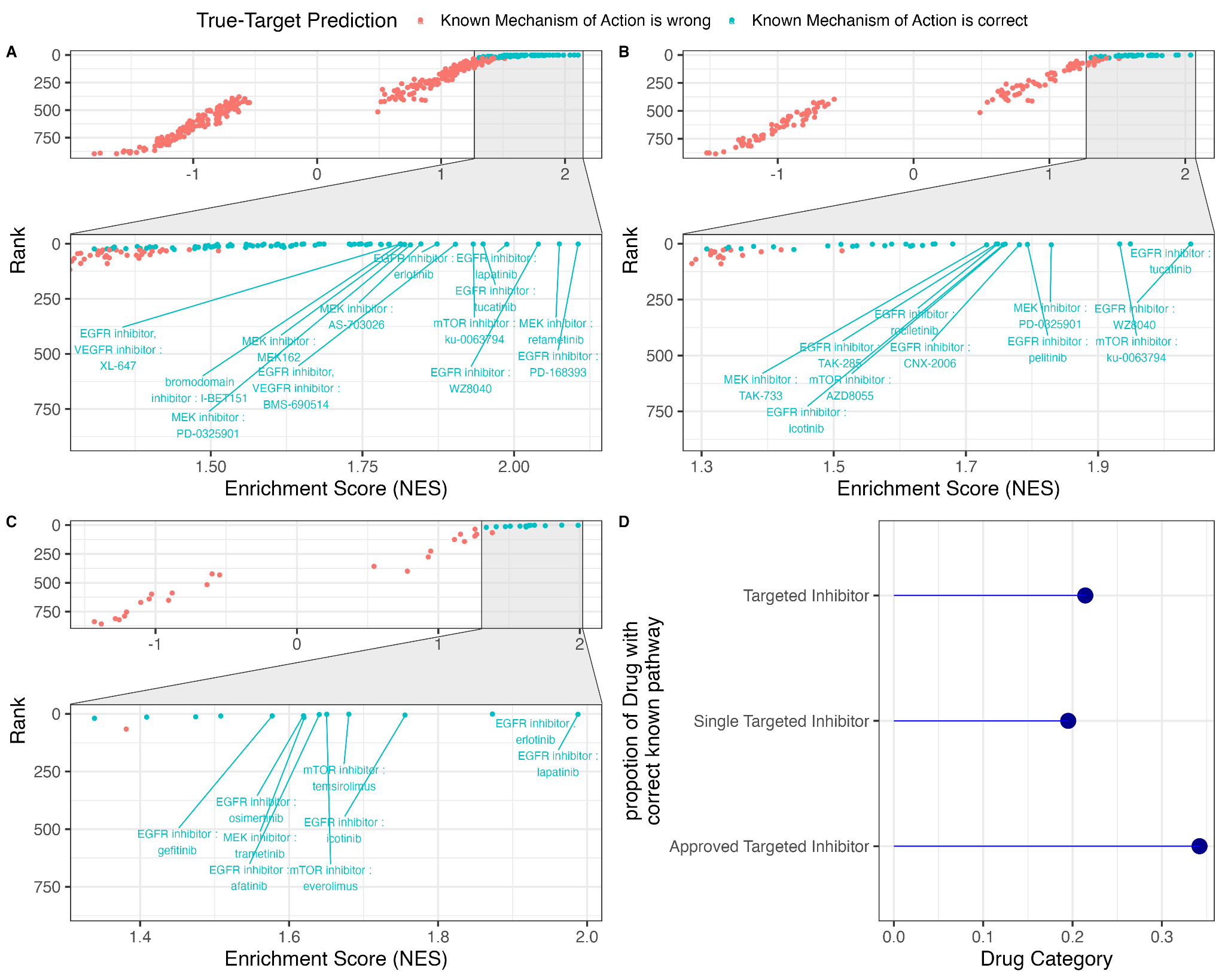

**Figure S10: Pathway-level annotation of the predicted mechanism of action.** Enrichment test **(A)** for drugs with targeted therapy and cancer inhibitors, where NES (Normalized Enrichment Score) is given on the x-axis and rank of the predicted rank of the known pathway is on the y-axis. The green color points represent the drug pathway which is predicted ranked < 27. Similarly, **(B)** Pathway enrichment test for drugs with single targets, and **(C)** only FDA-approved single-target cancer inhibitors. Drugs whose known targets are ranked within the predicted top 10 targets from DeepTarget are labeled in **(A, B, C)**. **(D)** We provided the proportion of drugs in each of these categories (Y-axis) that passes our threshold of Pathway rank< 27 (X-axis).

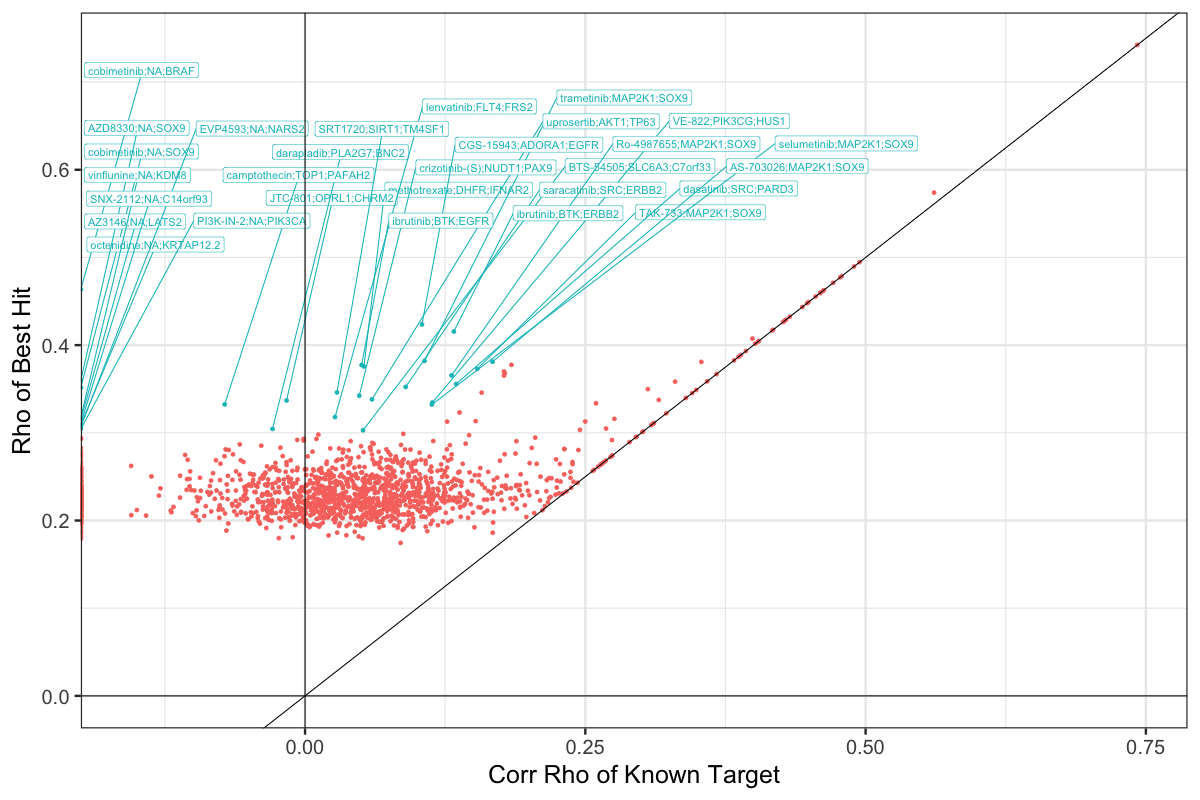

**Figure S11:** Correlation strength between drug response and CRISPR-KO of known target and the best-predicted target from DeepTarget.

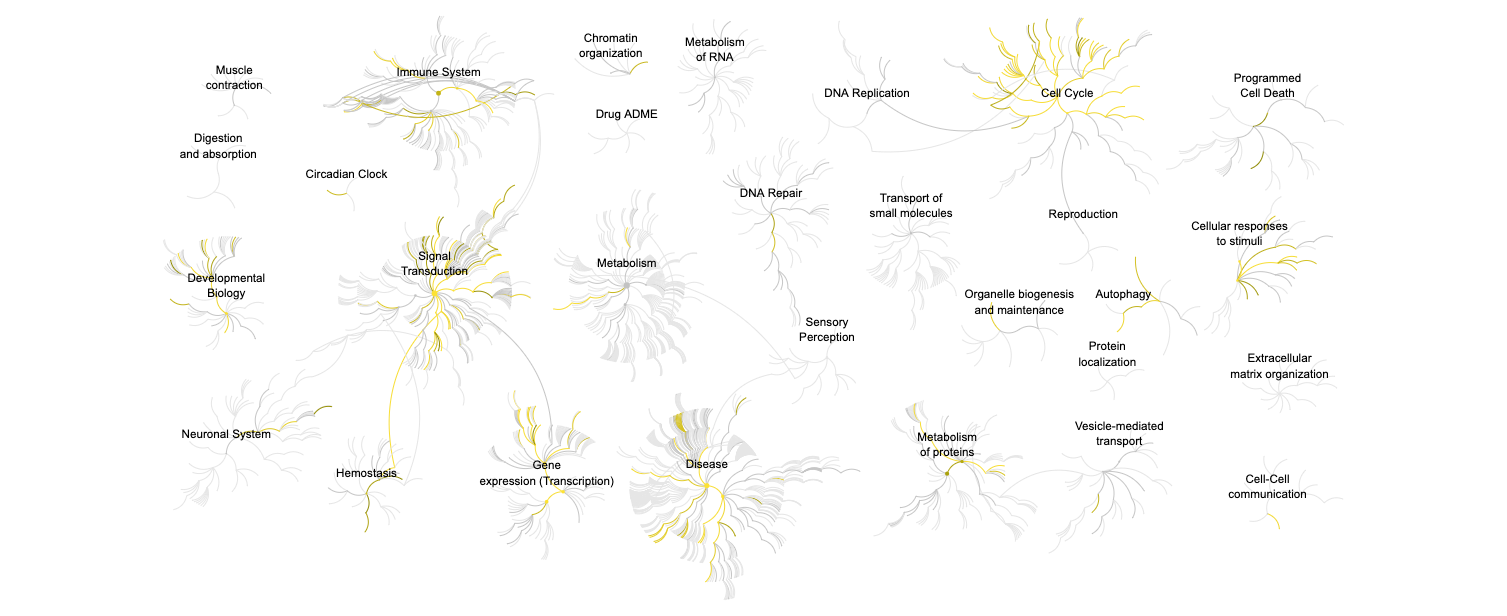

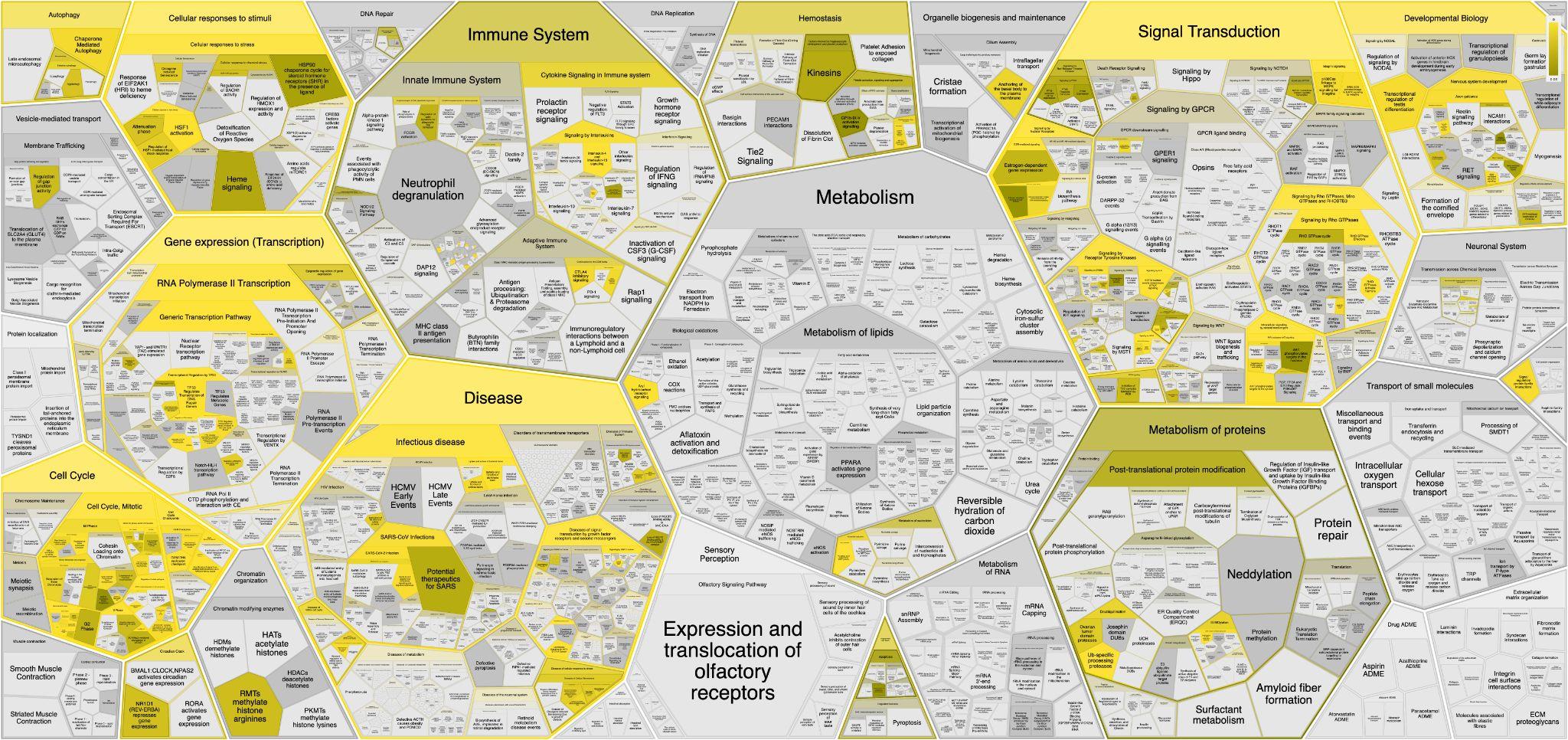

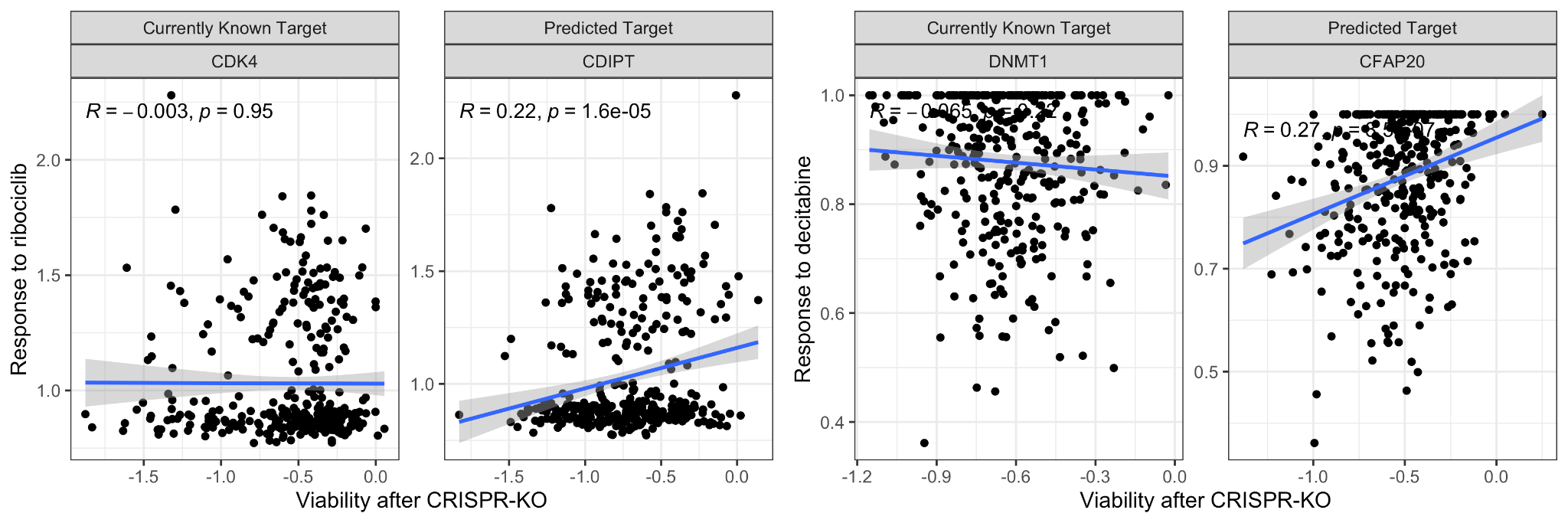

**Figure S12:** Pathways enriched in the mechanism of actions of drugs with incorrect targets **(A)** are shown at the Reactome level showing high-level pathways and **(B)** then at high-resolution at each subclass of pathways.

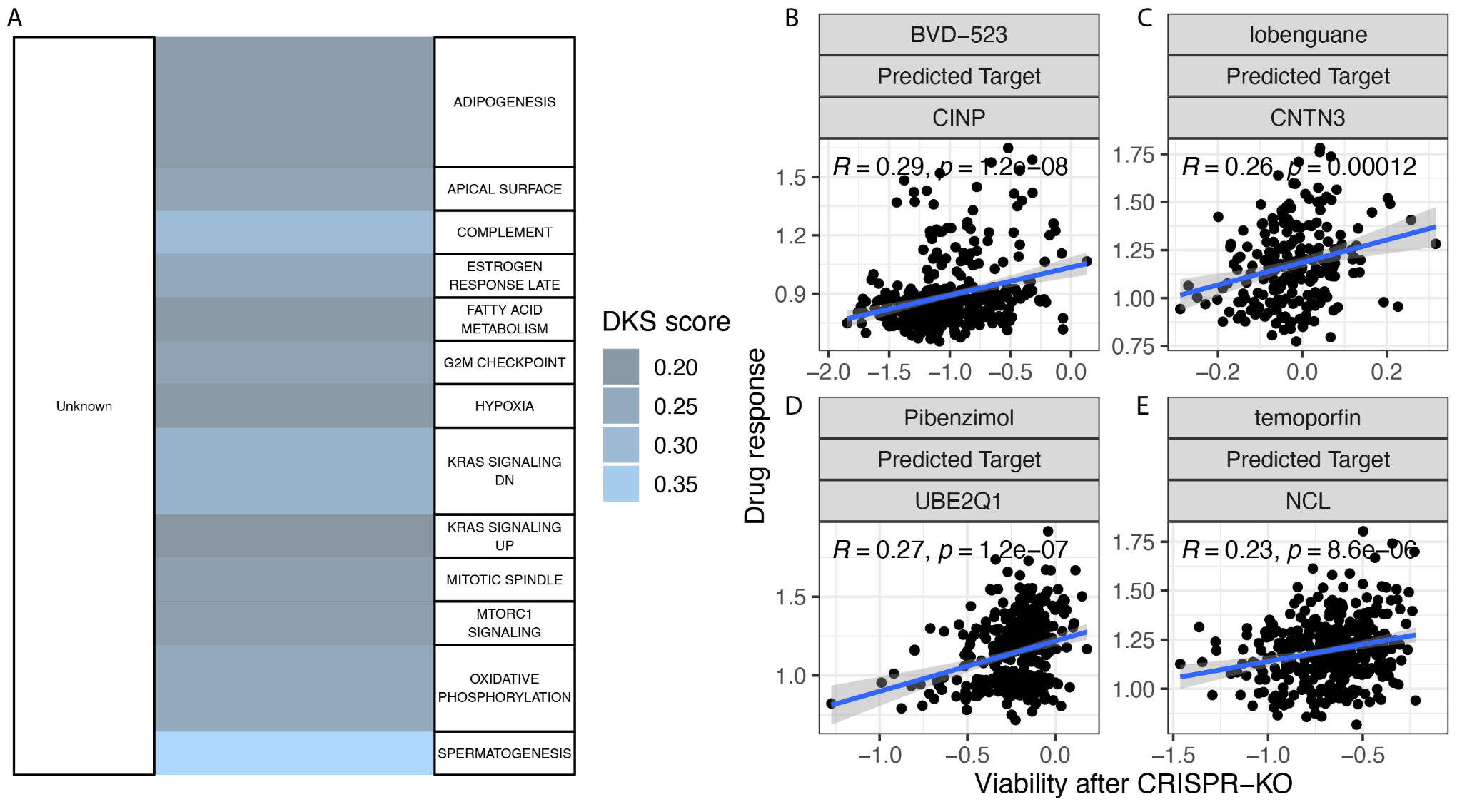

***Figure S13: Predicting new targets for cancer drugs with no known targets. (A)*** *A list of pathway annotations for the predicted target is provided in the right column where the left column represents the current status of the targets - Unknown. The intensity of the connection represents the predicted target DKS score and the width represents the number of drugs with such transition. The color legend is provided in right.* ***(B-D)*** *The correlation between viability after drug treatment (Y-axis) and after the knockout of the predicted target (X-axis) of the four top hits reported in the text. The drug name and the predicted target are provided in the top strips. The strength of correlation and regression line (blue) are provided.*

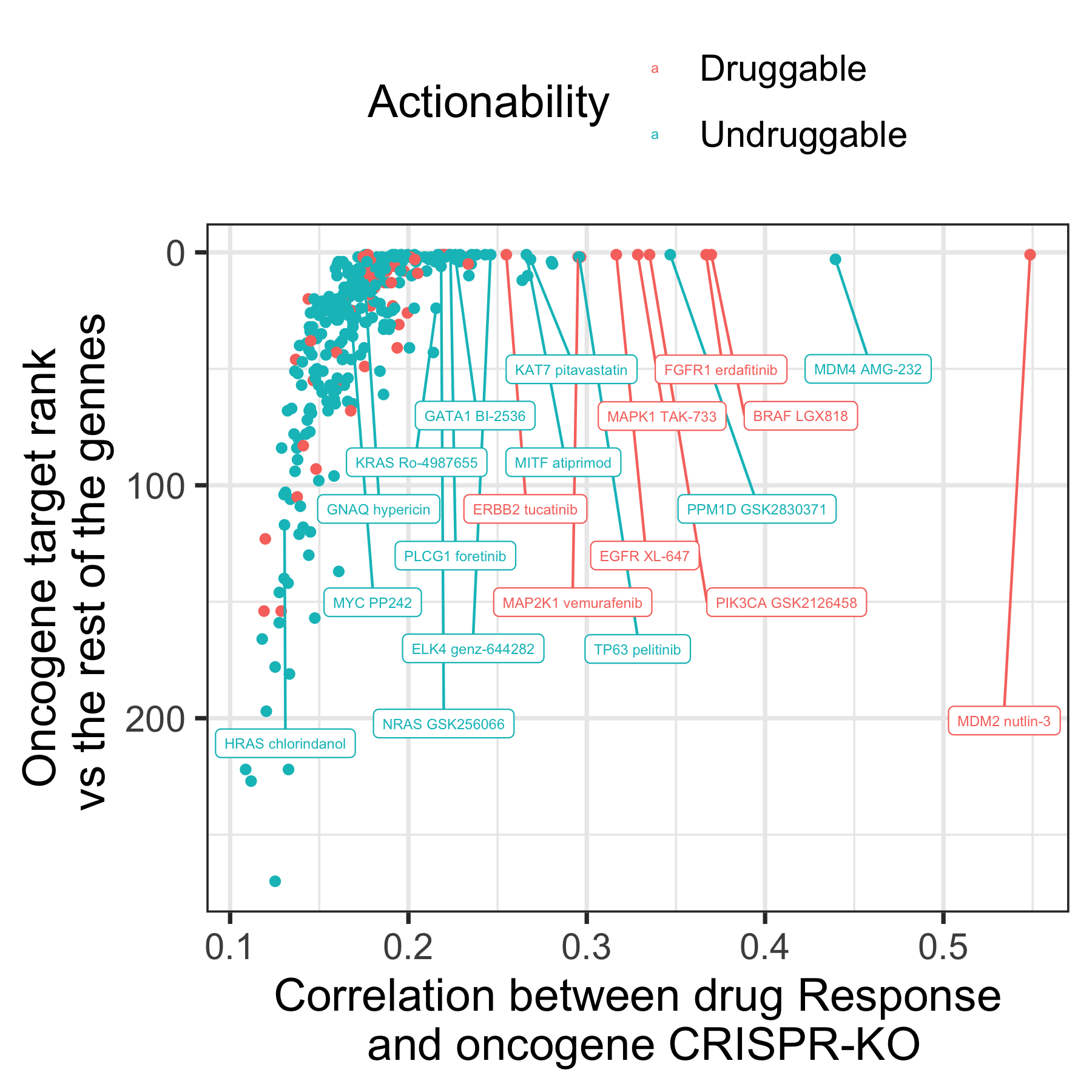

**Figure S14: Best targeting drug for all 250 oncogenes from COSMIC.**
